# Synchrony and idiosyncrasy in the gut microbiome of wild baboons

**DOI:** 10.1101/2021.11.24.469913

**Authors:** Johannes R. Björk, Mauna R. Dasari, Kim Roche, Laura Grieneisen, Trevor J. Gould, Jean-Christophe Grenier, Vania Yotova, Neil Gottel, David Jansen, Laurence R. Gesquiere, Jacob B. Gordon, Niki H. Learn, Tim L. Wango, Raphael S. Mututua, J. Kinyua Warutere, Long’ida Siodi, Sayan Mukherjee, Luis B. Barreiro, Susan C. Alberts, Jack A. Gilbert, Jenny Tung, Ran Blekhman, Elizabeth A. Archie

## Abstract

Human gut microbial dynamics are highly individualized, making it challenging to link microbiota to health and to design universal microbiome therapies. This individuality is typically attributed to variation in host genetics, diets, environments, and medications, but it could also emerge from fundamental ecological forces that shape microbiota more generally. Here we leverage extensive gut microbial time series from wild baboons—hosts who experience little interindividual dietary and environmental heterogeneity—to test whether gut microbial dynamics are synchronized across hosts or largely idiosyncratic. Despite their shared lifestyles, baboon microbiome dynamics were only weakly synchronized. The strongest synchrony occurred among baboons living in the same social group, likely because group members range over the same habitat and simultaneously encounter the same sources of food and water. However, this synchrony was modest compared to each host’s personalized dynamics. Indeed, host-specific factors, especially host identity, explained 10 times the deviance in longitudinal microbial dynamics, compared to factors shared across hosts. These results contribute to mounting evidence that highly idiosyncratic gut microbiomes are not an artifact of modern human environments, and that synchronizing forces in the gut microbiome (e.g., shared environments, diets, and microbial dispersal) are often not strong enough to overwhelm drivers of microbiome personalization, including host genetics, priority effects, horizontal gene transfer, and functional redundancy.

## Introduction

Mammalian gut microbiotas are highly complex, dynamic ecosystems. From these dynamics emerge a set of life-sustaining services for hosts, which help them digest food, process toxins, and resist invading pathogens. Despite their importance, our understanding of how gut microbial communities change over months and years within hosts, especially the collective dynamics of microbiotas from hosts living in the same population, is relatively poor^1, 2^. This gap exists in part because we lack time series data that track gut microbiota longitudinally across many hosts living together in the same population. As a result, it has been difficult to answer key questions. For example, when host populations encounter shifting environments and resources, does each host’s microbiota respond similarly—i.e., in synchrony—or idiosyncratically to these changes? Further, what factors predict synchronized versus idiosyncratic microbiota?

Answering these questions is important because synchronized gut microbial communities, if and when they occur^1^, could help explain shared microbiota-associated traits in host populations, such as patterns of disease susceptibility^3, 4^. A high degree of microbiome synchrony could also be good news for researchers working to predict microbial dynamics because it would suggest that similar ecological principles govern changes in microbial composition across hosts^5^. There is also theoretical justification to expect some degree of coordinated dynamics, as host populations and their microbiotas can be considered a ‘microbiome metacommunity’ (see e.g.,^6, 7, 8, 9^). Metacommunity theory predicts that synchrony will arise across microbiotas if their hosts experience similar environmental conditions and/or high rates of microbial dispersal between hosts^10, 11^. In support, fruit bats living in the same colony exhibit coordinated fur microbiota dynamics, and shared environments and microbial dispersal are both implicated in this synchrony^1^.

However, even in the presence of such synchronizing forces, there are many reasons to expect that hosts in a microbiome metacommunity will exhibit idiosyncratic (i.e., individualized) microbial compositions and dynamics. First, idiosyncratic dynamics are expected when the same microbes in different hosts respond in different ways to environmental fluctuations, chance events, and/or interactions with other microbes^12, 13, 14, 15^. These forces are likely to be important in gut microbiota where priority effects, functional redundancy, and horizontal gene flow can cause the same microbial taxon to perform different functions, play different ecological roles, and exhibit different environmental responses in different hosts (or cause different microbial taxa to play the same role in different hosts)^16, 17, 18, 19^. Second, several cross-sectional studies, in both humans and animals, find that individual hosts exhibit distinctive gut microbiota, and host identity explains a large fraction of population-wide microbiome taxonomic variation^1, 20, 21, 22, 23, 24, 25^. These results suggest that longitudinal gut microbial changes are also likely to be asynchronous across hosts. Indeed, some longitudinal studies in humans and animals find personalized gut microbial dynamics^1, 24, 26, 27, 28^, which are usually attributed to interpersonal differences in diet, medications, and lifestyle^27, 29, 30, 31^. If these explanations are correct, then observed idiosyncrasy in microbiome dynamics may be simply explained by a lack of shared environmental drivers rather than distinct microbiome responses to shared environments (but see ^27^). In contrast, if personalized dynamics persist even when hosts share the same environment, then (i) host-specific dynamics may not be solely attributable to interpersonal differences in lifestyles; (ii) predicting the dynamics of microbial taxa in individual hosts may prove difficult; and (iii) microbiome interventions that rely on manipulating taxa may face challenges beyond heterogeneity in lifestyles, and instead may be related to conserved ecological principles that govern the gut microbiome.

## Data and methods

Here we test the degree to which gut microbial taxonomic composition and dynamics, as measured via 16S rRNA gene sequencing, are synchronized versus idiosyncratic in wild baboon hosts living in the Amboseli ecosystem in Kenya^32^. Baboons are terrestrial primates that live in stable social groups, typically with 20 to 130 members. The 600 baboons in our data set lived in 12 social groups over a 14-year span (April 2000 to September 2013; 5 original groups and 7 groups that were fission/fusion products from these original groups; **Fig. 1A**). The baboons were members of the well-studied Amboseli baboon population^32^, which has been followed by the Amboseli Baboon Research Project since 1971. This project collected detailed longitudinal data on rainfall and temperature; social group membership, ranging patterns and diet; and host traits such as age, sex, social relationships, and dominance rank (see Supplementary Materials).

**Fig. 1.**
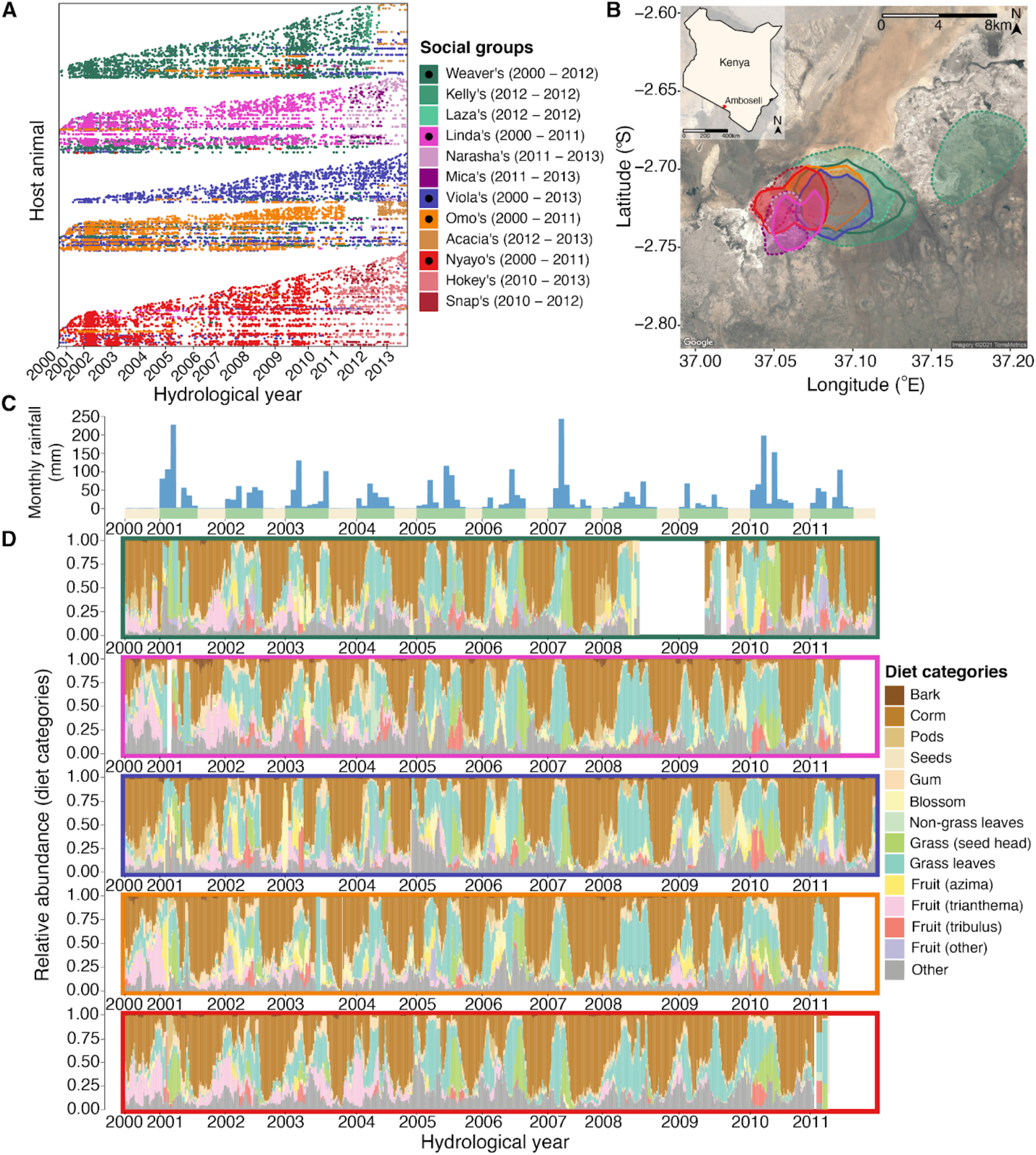
Baboons in Amboseli experience shared environments at multiple scales. **(A)** Our microbiota time series consisted of 17,265 16S rRNA gene sequencing gut microbial profiles. Each point represents a microbiota sample, plotted by the date it was collected (x-axis). Each row (y-axis) corresponds to a unique individual host. Samples were collected from 600 wild baboons living in 5 original social groups (indicated by dark colors marked with black dots in the legend) and 7 groups that fissioned/fused from these original groups (no black dots). **(B)** All baboon groups ranged over a shared ∼60 km^2^ area, and the social groups had largely overlapping home ranges. Ranges are shown as 90% kernel densities over the sampling period specific to each group; 5 original social groups are shown with solid borders, fission and fusion products with dashed borders. **(C)** Monthly rainfall amounts (blue bars, in mm) with yellow and green stripes along the x-axis representing dry and wet seasons, with the width of the green stripes reflecting the number of months within the focal year that had at least 1 mm rainfall. **(D)** Temporal shifts in diet from the years 2000 – 2013, shown as the relative abundance of diet components in the 5 original social groups over 30-day sliding windows prior to each sample collection date. Colors correspond to the 13 most common food types, while the grey bars correspond to other or unknown food types. Colored boxes around each panel reflect each of the 5 original, most extensively sampled social groups (colors as in plots A and B). The white bars indicate time periods where no diet data were collected.

Like many natural animal populations, the Amboseli baboons experience shared diets, environments, and opportunities for between-host microbial dispersal that have the potential to drive microbiota synchrony across hosts. Because baboons are not territorial, all 12 baboon social groups used an overlapping ∼60 km^2^ range (**Fig. 1B**; **video S1**;^33^). Hence all animals were exposed to similar microbes from the environment and shared seasonal changes in rainfall and temperature^33, 34, 35^. The Amboseli ecosystem is a semi-arid savanna with a 5-month long dry season spanning June to October, during which very little rain falls. The remaining 7 months (November to May) constitute the wet season, which is characterized by highly variable rainfall (**Fig. 1C**; mean annual rainfall between 2000 and 2013 was 319 mm; range = 140 mm to 559 mm). These seasonal shifts in climate drive a rotating set of foods consumed by the baboons: during the dry season the baboons rely largely on grass corms, shifting to growing grass blades and grass seed heads in the wet season (**Fig. 1D**). Within baboon social groups, diets and environments are especially congruent because group members travel together in a coordinated fashion across the landscape, encountering and consuming resources and feeding on the same seasonally available foods at the same time^33, 36, 37, 38, 39, 40^. Group members also groom each other, combing through each other’s fur and placing some items in their mouths, which may contribute to host-to-host microbial transmission^41^. Finally, at the level of individual hosts, host genetic variation has a consistent, albeit modest, effect on gut microbial composition in this population^33^. Other host-specific traits, like age, sex, and social status, also lead some individuals to share aspects of their behavior, immune profiles, and physiology, which could also lead to more congruent microbial dynamics.

A key advance in our study is longitudinal sampling of gut microbial composition from fecal samples collected from hundreds of known baboons throughout their lives (**Fig. 1A**). Such dense, long-term, longitudinal microbiota sampling is difficult to achieve in many animals, including humans. The 17,265 fecal samples in our study were collected from baboons who ranged in age from 7.4 months to 27.7 years, spanning these animals’ natural lifespans (**fig. S1A**). Each baboon was sampled a median of 19 times, and 124 baboons were sampled at least 50 times (**fig. S1B**). On average, these samples spanned 4.3 years of a baboon’s life (range = 4 days to 13.2 years; **fig. S1C**), with a median of 35 days between consecutive samples (**fig. S1D**).

A large majority of the microbiota data we use here were published in Grieneisen et al.^33^, but we include data from 1,031 additional samples that were generated at the same time using the same methods (they were not included in the heritability analysis of Grieneisen et al.^33^ because we lack pedigree information for these hosts). Briefly, we generated 896,911,162 sequencing reads (mean = 51,913.6 reads per sample; range = 1021 - 477,241, **fig. S1E**). We retained microbial amplicon sequence variants (ASVs) with a minimum of 3 reads per sample. DNA concentration and ASV diversity were not predicted by time since sample collection (**fig. S1G, S1H**). To allow us to compare the dynamics of individual taxa in different hosts, we focused on taxa that were common across hosts, retaining only those found in at least 20% of samples. This filtering resulting in 341 ASVs (mean = 162 ASVs per sample; range = 19 - 311 ASVs; **fig S1F; table S1**). While this filtering was somewhat stringent, it captured 92% of the reads and many of the same compositional properties of the data set when filtered to 5% prevalence (**fig. S2**). As is typical for wild microbiota, 22.9% of the 341 ASVs could not be assigned to a known family (78 of 341), and 5.5% of ASVs could not be assigned to a known phylum (19 of 341; **table S1**). To address the compositional nature of our data, read counts were centered log-ratio (clr) transformed independently in each sample (including independent transforms for samples from the same individual), prior to all analyses^42, 43^. Therefore, our results must be considered with respect to the chosen reference frame, which in this case is the geometric mean of taxon abundances in each sample, or the abundance of a sample’s ‘average microbe’.

To test whether shared environmental conditions and host traits lead to similar gut microbial compositions and synchronized dynamics across the microbiome metacommunity, we used three main approaches (see Supplementary Materials for details of all analyses). First, we characterized patterns of temporal autocorrelation in ASV-level Aitchison similarity within and between hosts over time. Our expectation was that, if hosts or social groups exhibit idiosyncratic composition and dynamics, then samples collected close in time from the same baboon, or from baboons in the same group, should be much more similar than they are to samples collected from different baboons living in different groups. Alternatively, if gut microbial dynamics are strongly synchronized, then samples collected close in time across the metacommunity should be compositionally similar, and samples collected from the same host should not be substantially more similar than samples from different baboons. These analyses were run in R (v 4.0.2; ^44^) using custom-written functions (code and analyzed data are available on GitHub/OSF; see Data Statement).

Second, to test whether dispersal limitation could explain microbiome idiosyncrasy, we estimated metacommunity-wide microbial migration probabilities in each season and year using the Sloan Neutral Community Model for Prokaryotes^45, 46^. This model assumes that each local community, defined as the ASV-level microbial composition of a single host in a given season-year combination, is the outcome of stochastic population dynamics and microbial immigration from other hosts in the microbiome metacommunity (i.e., other local communities). Briefly, local communities have a constant size *N*, and individual microbes within each local community die at a constant rate. These deaths create vacancies that can be occupied, either by individuals immigrating from the microbiome metacommunity (with probability *m*), or by daughter cells from any taxon within the local community (i.e., from reproduction within the same host, with probability 1-*m*). Taxa that are common in the metacommunity have a higher chance of occupying vacancies than rare taxa. Without immigration from the microbiome metacommunity, ecological drift leads each host’s microbial diversity to reduce to a single taxon. Thus, the migration probability, *m*, represents the metacommunity-wide probability that any taxon, randomly lost from a given host/local community, will be replaced by dispersal from the microbiome metacommunity, as opposed to reproduction within hosts^45, 46^. Following Burns et al.^47^, *m* can be interpreted as a measure of dispersal limitation, such that low migration probabilities signify high dispersal limitation. We estimated season and hydrological year-specific values for *m* by defining the microbiome metacommunity as either the hosts’ social group or the whole host population. We fit neutral models using nonlinear least-squares regression as implemented in the R package tyRa^48^.

Third, to quantify the relative magnitude of idiosyncratic versus synchronized dynamics for community metrics and common families and phyla, we used generalized additive models (GAMs) to capture the non-linear, longitudinal dynamics of 52 features, including the first three principal components of ASV-level composition, three indices of alpha diversity (ASV richness, the exponent of ASV-level Shannon’s H, and the inverse Simpson index for ASVs, as computed by the function reyni from the R package vegan^49^), and the clr-transformed abundances of 12 phyla and 34 families present in >20% of samples. We analyzed phyla and families (as opposed to genera or ASVs) because phyla and families are highly prevalent across samples (mean prevalence = 85.6% for the 12 phyla and 73.7% for the 34 families), offering excellent power to compare their dynamics between different baboons. We note, however, that phyla and families might be expected to exhibit stronger synchrony than lower-level taxa because, compared to species or strains, the dynamics of families and phyla reflect multiple microbial processes and interactions, which are expected to buffer them against large fluctuations in abundance. Further, the processes and interactions that a given phylum or family collectively encompasses may be more consistent across hosts than those carried out by an individual species or strain (although this consistency will vary depending on the phylum, family, or process in question^18, 50^).

Our GAMs allowed us to calculate the percent deviance in each feature’s dynamics attributable to factors that could contribute to synchronized dynamics at different scales (percent deviance is a measure of goodness-of-fit for nonlinear models and is analogous to the unadjusted R^2^ for linear models). We considered deviance explained by factors at three scales: those experienced by the whole host population (e.g., rainfall and temperature), those differentiated by social groups (e.g., group identity, group home range location, and diet), and those differentiated at the level of individual hosts (e.g., host identity, sex, age, and social dominance rank; see below for complete model structures). If microbiome community dynamics are largely idiosyncratic, then population- and group-level factors will not explain considerable deviance in microbiota change over time, and instead, a large fraction of the deviance will be attributable to host identity, controlling for shared environments, behaviors, and traits. Alternatively, if shared environments and behaviors across the population and within social groups synchronize gut microbiota, then population- and group-level factors should explain substantial deviance in community dynamics. To ensure sufficiently dense sampling for identifying host- and group-level dynamics, all three GAMs were run on a subset of the full data set, consisting of 4,277 16S rRNA gene sequencing profiles from the 56 best-sampled baboons living in the 5 social groups sampled the longest (between 2002 and 2010; median = 72.5 samples per host; minimum = 48 samples; maximum = 164 samples; **fig. S3**). GAMs were fit using the R package mgcv^51, 52, 53^.

Notably, the GAM approach allows us to identify the percent deviance attributable to host identity, but does not identify the specific characteristics that account for host identity effects. Genetic effects are a likely candidate, as previous analyses demonstrate that taxon abundance and summaries of gut microbiome position are lowly to moderately heritable in this population^33^. To evaluate this possibility, we performed a *post hoc* analysis of the relationship between the deviance explained in the GAMs for each microbial taxon and the heritability of that taxon’s relative abundance^33^. If host effects on microbiome dynamics are in part explained by host genotype, we predicted that taxon heritability should be positively correlated with deviance explained at the host level (i.e., model P+G+H), but not at the group or population level (i.e., model P and model P+G).

## Results and Discussion

### Baboon gut microbiota exhibit cyclical shifts in community composition across seasons and years

We began by visualizing annual and inter-annual fluctuations across all 600 hosts (17,265 samples) over the 14-year span of the data. Consistent with prior research on primates^54, 55, 56^, we found population-wide, cyclical shifts in microbiome community composition across seasons and years (**Fig. 2**). This wet-dry seasonal cyclicity was primarily observable in the first principal component (PC1) of a principal component analysis (PCA) of clr-transformed ASV read counts (**Fig. 2A, 2B**; **fig. S4-S6**; PC1 explains 16.5% of the variance in microbiome community composition). PC1 exhibited its lowest values during the dry season, and highest values during the wet season, mirroring monthly rainfall (**Fig. 2B; fig. S6**). Consistent with these annual, seasonal fluctuations, periodicity for PC1, calculated as the mean distance between major peaks, was 374 days. PC1 was also linked to annual rainfall across years, exhibiting especially low values throughout 2008 and 2009, which corresponded to the worst continuous drought in the Amboseli ecosystem in 50 years (**Fig. 2A, 2B**). We also observed small, but statistically significant seasonal differences in PC2 and PC3 (8.4% and 3.7% of variation in community composition; **Fig. 2C**; **fig. S4-S6**) and in measures of alpha diversity (**Fig. 2C**; **fig. S6, S7**), as has been reported in other ecosystems^57^. Together, these seasonal changes are probably caused by seasonal shifts in plant phenology and its effects on diet (**Fig. 1D**), as well as the effects of rainfall and other weather variables on bacterial exposures from the abiotic environment (e.g., soil communities and sources of drinking water).

**Fig. 2.**
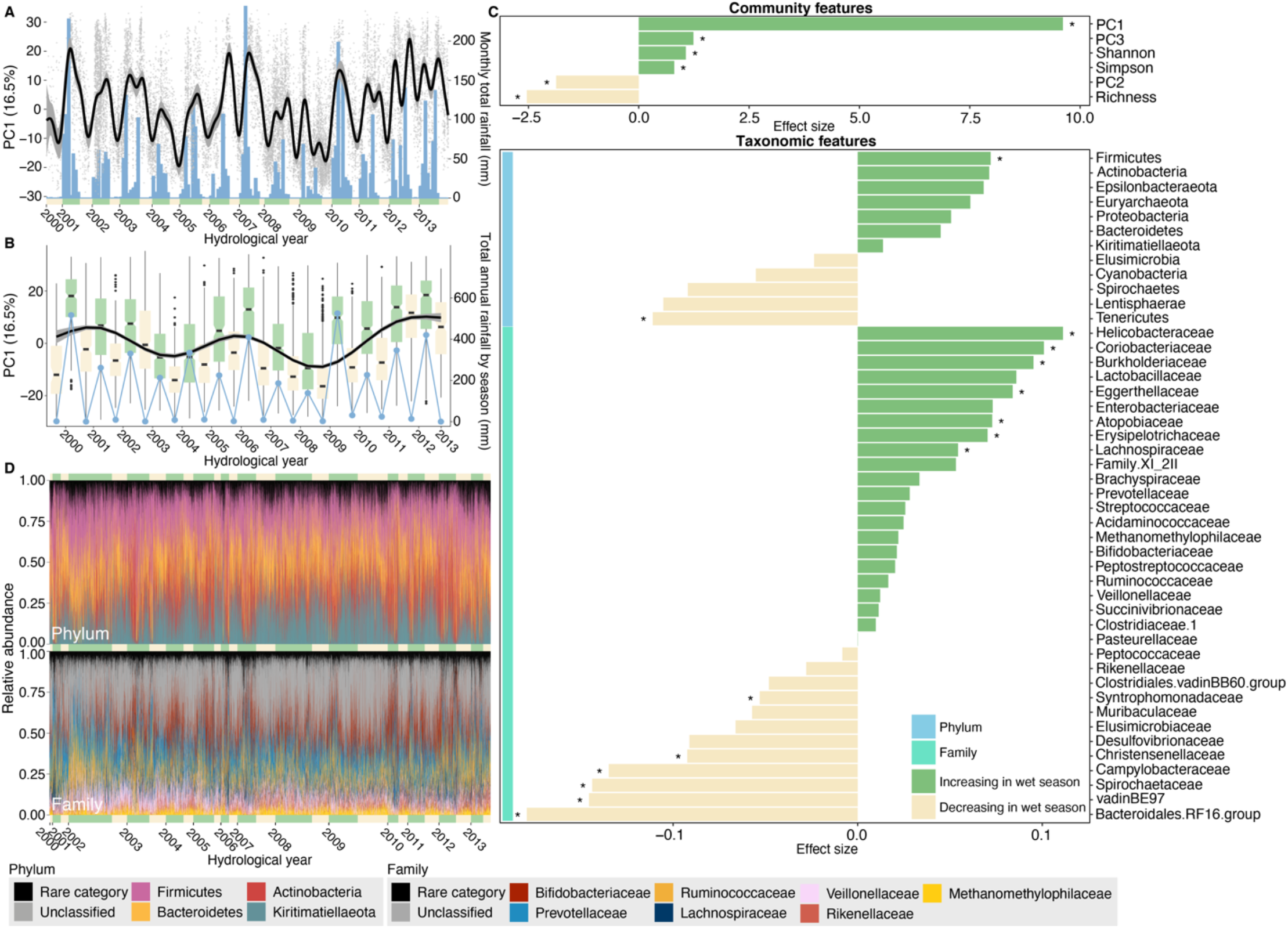
Baboons show population-wide, cyclical shifts in microbiome community composition across seasons and years. **(A)** Changes in microbiome PC1 mirror monthly rainfall across all 14 years. The grey points show values of PC1 for each of the 17,265 samples (y-axis) on the dates they were collected (x-axis). The black line and its grey band show the predicted daily trend for PC1 across samples and its 95% simultaneous confidence interval, treating time as a continuous variable from April 21, 2000 to September 19, 2013. Blue bars show monthly rainfall (right-hand y-axis), and the yellow and green bars along the x-axis represent dry and wet seasons, respectively, with the width reflecting the number of months within the focal year with at least 1 mm rainfall. **(B)** Changes in microbiome PC1 on an annual scale across all 14 years (N = 17,265 samples). The box plots show the distribution of PC1 in wet (green) and dry (yellow) seasons. The black line shows the estimated annual trend for PC1 across all hydrological years, and the blue points show total annual rainfall (right-hand y-axis). **(C)** The effect of season varies across 52 features of the microbiome, including six community features (top panel) and 46 taxa (bottom panel; 12 phyla: light blue vertical bar; 34 families: turquoise vertical bar; for 341 ASVs, see **fig. S13**). Each horizontal bar shows the effect of season from linear mixed models for each feature. Asterisks indicate features that changed significantly between the wet and dry seasons (N = 17,265 samples; FDR threshold = 0.05). See **figs. S8, S9** for feature-specific smooths and **fig. S10** and **table S3** for results for ASVs. **(D)** Bar plots showing the relative abundance of ASVs colored by four most common microbial phyla (above) and the seven most common families (below) across all 17,265 samples. Green and yellow bars along the x-axes represent wet and dry seasons, with the width corresponding to the number of samples in the focal hydrological year and season. 22.9% of ASVs (78 of 341) could not be assigned to a known family (“unclassified”, shown in grey). The abundance of ASVs unclassified to family in the lower plot is ∼35% because one unclassified ASV was the second most abundant ASV in the data set, with a mean abundance of 16.9% across all samples (ASV#2, phylum Kiritimatiellaeota, order WCHB1-41; **table S1**).

In terms of individual microbiome taxa, 17% of phyla (2 of 12) and 38% of families (13 of 34) exhibited significant changes in relative abundance between the wet and dry seasons (**Fig. 2C**; **table S2**; linear models with false discovery rate (FDR) threshold = 0.05). These changes were significant for the phyla Firmicutes and Tenericutes (**Fig. 2C, 2D**; **fig. S8**), and were especially pronounced for the families Helicobacteraceae, Coriobacteriaceae, Burkholderaceae, Bacteroidales RF16 group, vadinBE97, Spirochaetaceae, and Campylobacteraceae (**Fig. 2C**; **fig. S9**). 28% of ASVs also exhibited significant changes in abundance across seasons (97 of 341 ASVs; linear models with FDR threshold = 0.05 for n = 393 models; **fig. S10**; **table S3**). However, most gut microbial ASVs, families and phyla did not exhibit significant changes in abundance across seasons, suggesting that many taxa play consistent roles in the gut throughout the year, including Kiritimatiellaeota, Elusomicrobia, Ruminococcacaceae, Clostridiaceae 1, and Rikenellaceae (**Fig. 2C**; **fig. S8, S9**; **table S2**).

### Baboon gut microbiota exhibit stronger idiosyncrasy than synchrony

While the microbiome metacommunity exhibited cyclical, seasonal shifts in composition, microbiome dynamics across different baboons were only weakly synchronized. Instead, consistent with prior observations of microbiome personalization^1, 20, 21, 22, 23, 24, 25^, patterns of temporal autocorrelation indicated that each baboon exhibited largely individualized gut microbiome compositions and dynamics (**Fig. 3**). In support, ASV-level Aitchison similarity was much higher for samples collected from the same baboon within a few days of each other than for samples from different baboons over the same time span, regardless of whether those animals lived in the same or a different social group (**Fig 3A, 3B**; Kruskal-Wallis: p < 2.2×10^−16^ for all comparisons). Likewise, a PERMANOVA of Aitchison similarities between 4,277 samples from the 56 best-sampled hosts revealed that host identity explained 8.6% (p < 0.001) of the variation in community composition, much larger than sampling day or month (R^2^ = 2.5% and 1.4%), group membership (2.2%), or the first three principal components of diet (0.04% to 2.4%; **table S4**; **fig. S11**).

**Fig. 3.**
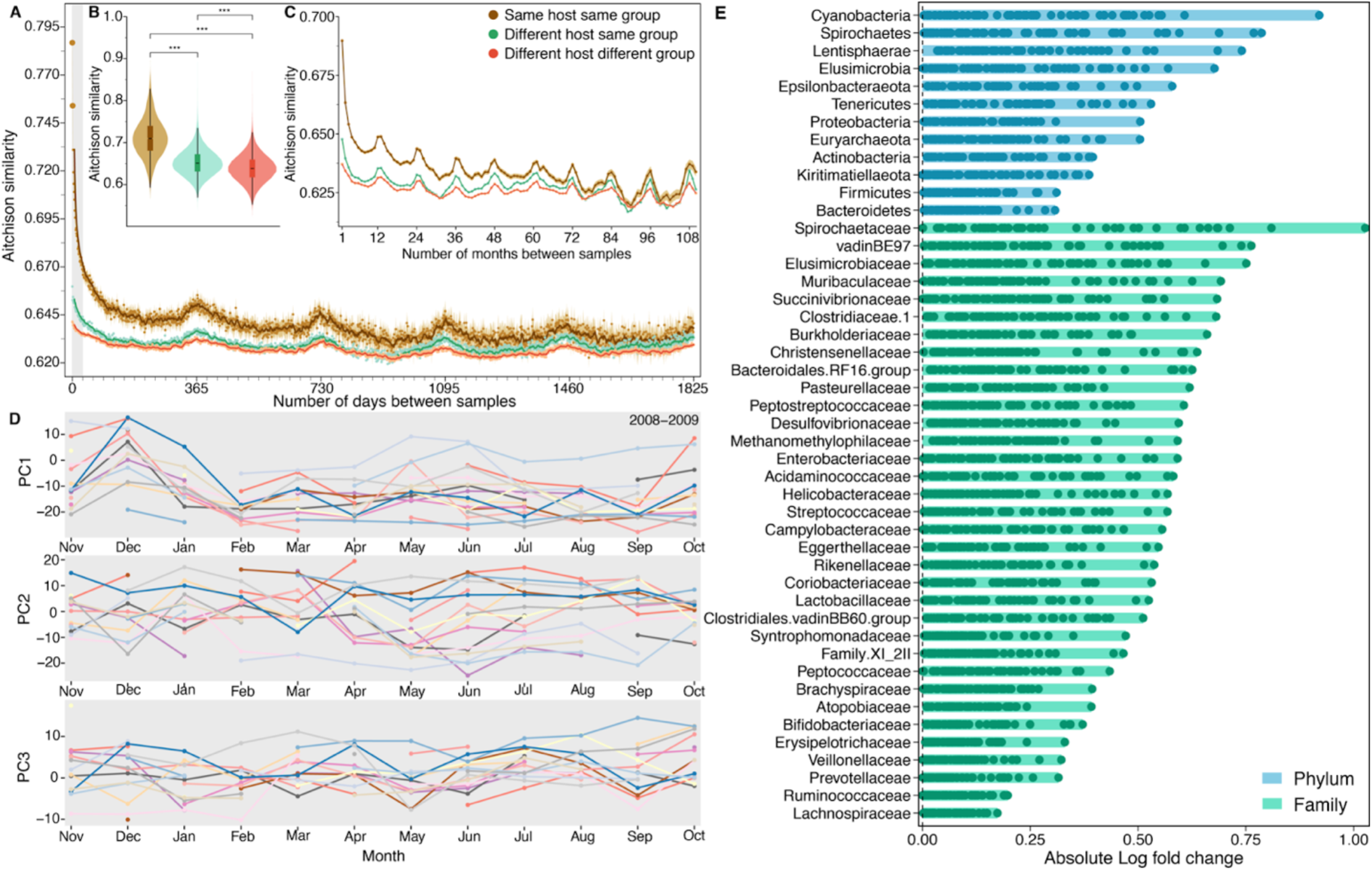
Baboons exhibit largely idiosyncratic gut microbial compositions and dynamics. **(A)** Temporal autocorrelation in baboon gut microbiome communities for samples collected on the same day and up to 5-years (1,825 days) apart. Points show mean ASV-level Aitchison similarity (y-axis) between samples as a function of the number of days between sample collection (x-axis; small tick marks correspond to months). Lines depict moving averages (window size = 7 days). The grey region on the left indicates samples collected within one month of each other. Brown points show average Aitchison similarity between samples collected from the same baboon (N = 392,817 distinct sample pairs from 547 hosts with 2 or more samples); green points show similarity between samples from different baboons living in the same social group (N = 16,391,761 distinct sample pairs); orange points show similarity between samples from different baboons living in different social groups (N = 77,520,289 distinct sample pairs). **(B)** Average Aitchison similarity between pairs of samples collected within 10 days of each other. Samples from the same baboon are significantly more similar than samples collected from different baboons in the same or different social groups (Kruskal-Wallis; p = 2.22 × 10^−16^; N distinct sample pairs = 5,791 for within-host comparisons; 218,340 for different host same group; 779,054 for different host different group). **(C)** Temporal autocorrelation in Aitchison similarity on monthly scales for samples collected up to 10 years apart (N distinct sample pairs = 496,057 for within-host comparisons; 23,433,667 for different host same group; 114,170,919 for different host different group). **(D)** Microbiome dynamics for 174 samples from 17 baboons for which we had at least one sample from 10 or more months during the 2008-2009 hydrological year (Nov 2008 to Oct 2009). Panels show each individual’s mean values for microbiome PC1, PC2, and PC3; each colored line represents a distinct host. See **fig. S14** for similar results during another densely sampled time period. Gaps indicate that the focal host did not have a sample in a given month. **(E)** Some taxa have more idiosyncratic abundances than others. Each horizontal bar shows a given taxon’s minimum and maximum absolute log fold change in abundance across the 56 best-sampled hosts (hosts are represented as points within the bars; see **fig. S3** for information on the 4,277 samples from the 56 best-sampled hosts). Absolute fold changes were calculated, for a given taxon in a given host, as the taxon’s average clr-transformed abundance across all samples from that host, relative to the taxon’s grand mean in all hosts in the population. Hosts with large absolute fold changes for a given taxon therefore have abundances of that taxon that are either well above or below-average compared to its abundance in the host population at large (hosts with points close to zero exhibited taxonomic abundances typical of the population at large). For many taxa, hosts varied in their absolute log ratio values, indicating that they also deviated substantially from each other in the abundance of those taxa. Taxa (y-axis) are ordered (from top to bottom) by their highest absolute log ratio value across the 56 best-sampled hosts. Blue bars represent microbial phyla; green bars represent families. See **fig. S15** for a longitudinal version of this analysis for the most and least idiosyncratic phyla and families.

Aitchison similarity among samples from the same baboon fell steeply for samples collected a few days to a few months apart, indicating that individualized dynamics are strongest for samples collected close in time (**Fig. 3A-3C)**. At longer time scales (e.g., months and years), self-similarity was modest, but samples from the same baboon were significantly more similar to each other than they were to samples from different baboons, even for samples collected nearly three years apart (**Fig. 3A, 3C**; **fig. S12**). Following the initial steep decline in self-similarity, community similarity rose again slightly at 12-month intervals, both within and between hosts, reflecting synchronized, seasonal microbial dynamics across the host population. These small, 12-month peaks in similarity were visible even for samples collected more than 5 years apart, indicating that individual hosts and the population at large return to somewhat similar microbiome community states on 12-month cycles over several years (**Fig. 3C**). Hence, the patterns in **Fig. 3A** and **3C** show both idiosyncratic and synchronized microbial dynamics: over short time scales, hosts are much more similar to themselves than they are to others, but on annual scales, all hosts are weakly synchronized across seasons.

The greater influence of individualized dynamics compared to synchronized dynamics can also be captured by comparing microbiome dynamics for deeply sampled hosts sharing the same habitat at the same time (**Fig. 3D**; **fig. S13**). For instance, during the 2008-2009 hydrological year, we were able to collect nearly one sample per month from 17 individual baboons. When we aligned these time series, we observed little evidence of shared changes in the top three principal components of ASV-level microbiome composition across time, beyond some overall seasonal patterns in PC1. We also observed little convergence to similar values within any given month (**Fig. 3D**). Consequently, the microbiome of each baboon took a different path over the ordination space over the same 1-year span (**fig. S13**). We found similar results for another dense sampling period in the 2007-2008 hydrological year (**fig. S14**).

Microbiome taxa varied in their contributions to individualized gut microbiome compositions (**Fig. 3E**; **fig. S15**). For example, for the 56 best-sampled hosts (**fig. S3**), several phyla and families exhibited substantial variation in host mean clr-transformed abundance (i.e., across repeated samples for that host) compared to their mean clr-transformed abundance across all hosts. These taxa included members of the phyla Cyanobacteria, Spirochaetes, Lentisphaerae, and Elusimicrobia, and the families Spirochaetaceae, vadinBE97, Elusimicrobaceae, and Muribaculaceae (**Fig. 3E**; **fig. S15**). These highly variable taxa tended to exhibit, on average, below-average abundance compared to less variable taxa, which tended to exhibit, on average, above-average abundance. Thus, idiosyncratic dynamics may be more often linked to uncommon than common taxa, perhaps because uncommon taxa have greater functional variability across hosts (**fig. S16**).

To test whether individualized gut microbiome compositions and dynamics could be explained by microbial dispersal limitation between hosts, we used the Sloan Neutral Community Model for Prokaryotes to estimate metacommunity-wide migration probabilities, *m*, for ASVs in each season and hydrological year ^45, 46^. As described above, *m* provides a measure of dispersal limitation because it represents the probability that “vacancies” in a local community (i.e. a host’s microbiome) will be replaced by the process of dispersal from the microbiome metacommunity (i.e. other hosts), as opposed to reproduction within a focal host’s microbial community ^45, 46^. We found little evidence that dispersal limitation contributed to idiosyncratic compositions and dynamics. The estimated probability that a given ASV lost from a host’s microbiota would be replaced by an ASV from another host in the population was nearly 40% (the average host population-wide *m* across season and hydrological years = 0.373; range = 0.332 to 0.416; black points on **fig. S17**). These migration probabilities are generally lower than those Sieber et al. ^9^ found for marine sponges sampled from the same coastal location (range of *m* across sponge species: min=0.36; median=0.78; max=0.86) but much higher than for mice and nematodes, both in natural and laboratory populations (mice: *m*_wild_ = 0.11 and *m*_lab_ = 0.18; nematode: *m*_wild_ = 0.03 and *m*_lab_ = 0.01). Thus, they indicate that dispersal limitation is low for baboon microbiota in Amboseli.

Interestingly, when we re-defined the microbiome metacommunity to be the host’s social group, instead of the whole host population, migration probabilities were similar (average *m* across groups = 0.355; range = 0.347 to 0.365; colored points on **fig. S17**). Hence, despite several studies that find microbiome compositional differences between hosts living in different social groups, including in the Amboseli baboons^1, 41, 58, 59, 60, 61^, social group membership does not represent a major barrier to microbial colonization between baboons, perhaps because of their overlapping home ranges, similar diets, and network connections via male dispersal (e.g., **Fig. 1**).

### Shared environmental conditions are linked to modest synchrony across hosts

To quantify the relative magnitude of idiosyncratic versus synchronized gut microbial dynamics across the host population, social groups, and individual hosts, and to test whether synchrony varies for a set of common microbial taxa, we used generalized additive models (GAMs) to capture the nonlinear, longitudinal changes in 52 microbiome features (3 PCs of ASV-level community variation, 3 metrics of ASV-level alpha diversity, and clr-transformed relative abundances of 12 phyla and 34 families). For each feature, we ran three GAMs to measure the deviance explained in gut microbial dynamics by successive sets of parameters, reflecting the nested nature of our variables (**Fig. 4A**; x-axis of **Fig. 4C**; **table S5**). The population-level model (i.e., model P) captured factors experienced by the whole host population, including average rainfall and maximum daily temperature in the 30 days before sample collection and random effect splines to capture monthly and annual cyclicity in microbiome features (e.g., **Fig. 2A and B**; see **fig. S18** for effects of time of day, which was not included in the model). The group-level model (i.e., model P+G) included all the predictor variables in model P, and added a random effect spline for each social group, as well as variables to capture temporal changes in each group’s diet, home range use, and group size (**Fig. 4A, 4C**). The host-level model (i.e., model P+G+H) included all of the predictor variables in model P+G, and added a random effect spline for each host, and variables for host traits, including sex, age, and social dominance rank (**Fig. 4A, 4C**).

**Fig. 4.**
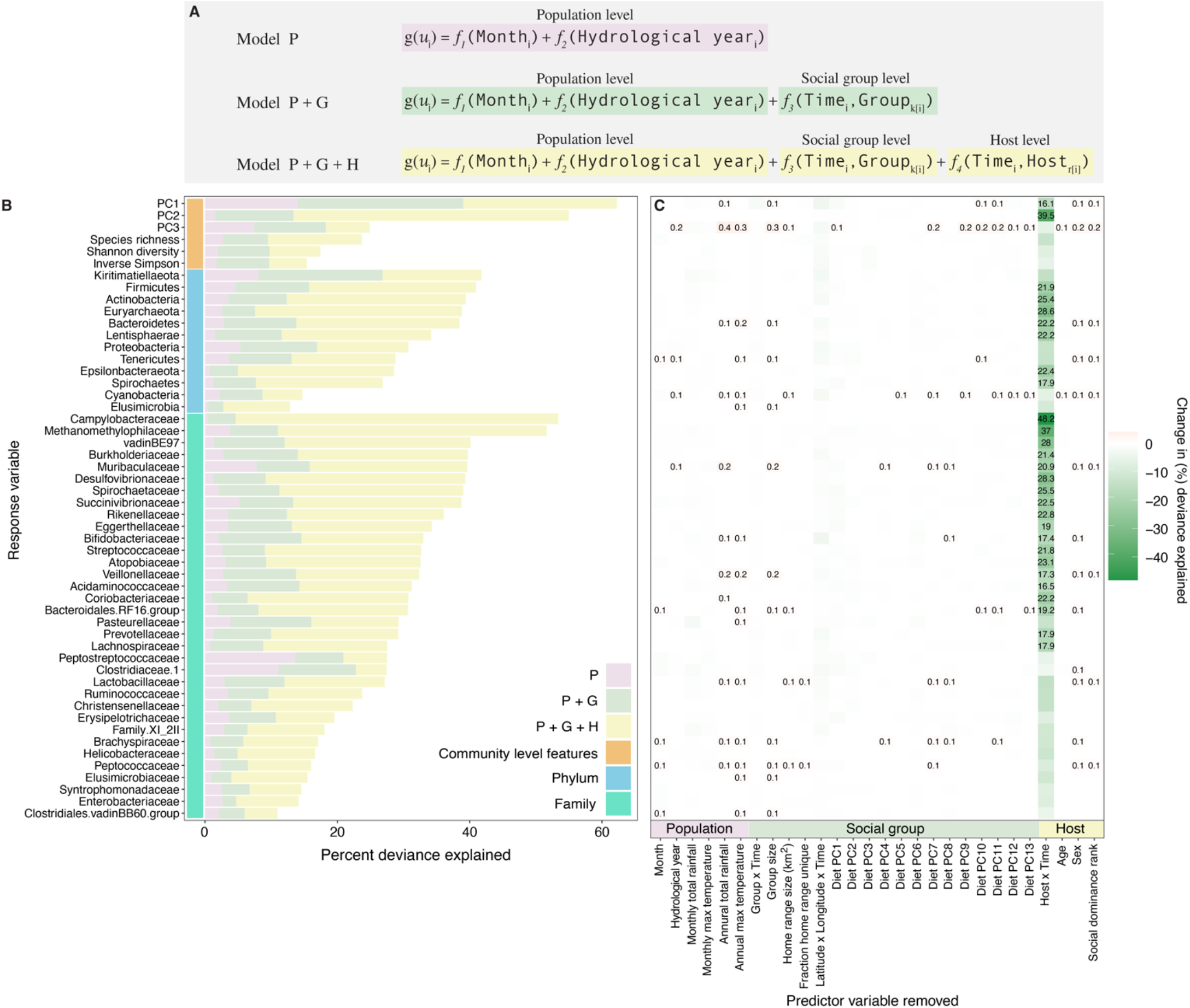
Multilevel modeling identifies idiosyncratic microbial dynamics. (**A**) We fit three hierarchical GAMs to 52 microbiome features measured in 4,277 samples from the 56 best-sampled baboons, all of whom lived in the 5 social groups sampled the longest (between 2002 and 2010; median = 72.5 samples per host; minimum =48 samples; maximum = 164 samples; **fig. S3**). Each model contained successive sets of predictor variables reflecting population-level factors (pink), group-level factors (green) and host-level factors (yellow). The factors at each level are listed at the bottom of panel C and defined in **table S5**). Panel (**B**) shows, for each microbiome feature (i.e., response variable), the deviance explained by model P and the successive sets of predictor variables added in model P+G and model P+G+H, respectively (**table S6**; percent deviance is a measure of goodness-of-fit for nonlinear models and is analogous to the unadjusted R^2^ for linear models). Panel (**C**) shows the loss in deviance explained for model P+G+H as we successively removed each predictor variable in turn from model P+G+H, keeping the model otherwise intact (**table S7**). Losses in deviance are shown in green, and we only provide numeric values for losses in deviance > 15%. Gains in deviance are shown in red; we only show numeric values for gains > 0.1%.

Consistent with our autocorrelation analyses (**Fig. 3**), comparing the deviance explained for each microbiome feature across the three models revealed stronger idiosyncratic than synchronized dynamics for most microbiome features (**Fig. 4B, 4C**). Indeed, host-specific factors, especially host identity, explained, on average, 10 times the deviance in the longitudinal dynamics of microbiome features, compared to factors shared across hosts (i.e., group-level or population-level factors). Specifically, model P only explained on average 3.3% (range = 0.46% to 14.0%) of the deviance across all 52 microbiome features (pink bars in **Fig 4B**; **table S6**), compared to 8.1% on average after adding group-level factors to the population-level model (increase from model P to model P+G; range = 2% to 25%; green bars in **Fig. 4B**; **table S6**), and 30.1% of the deviance after including host-level dynamics (model P+G+H; range = 11.0% to 62.2%) for the same set of features (yellow bars in **Fig. 4B**; **table S6**). Importantly, the added deviance for model P+G+H compared to model P or model P+G was not simply caused by including more parameters. Specifically, randomizing host identity and host-level traits across samples, while keeping each sample’s annual, seasonal, and group identity intact, led to a substantial drop in deviance explained compared to the real data (**fig. S19**). For instance, for PC2, which was most strongly associated with host-level effects among all three PCs, the deviance explained by model P+G+H dropped from 55% to 16.6% when host identity and traits were randomized (**fig. S19;** see supplement and **fig. S20** for an additional analysis investigating the effect of model complexity on deviance explained). That said, for PC3, randomized host-level dynamics resulted in a 3% increase in deviance explained compared to the P+G model. While less than the increase observed in the real data (6.6%), this result suggests that deviance explained by the P+G+H model may be modestly inflated for some microbiome features.

44 of the 52 microbiome features exhibited greater gains in deviance explained by adding host-level factors to model P+G, compared to adding group-level factors to model P. Of these 44, 22 features gained more than 20% deviance explained between model P+G and model P+G+H (**Fig. 4B**; **table S6**). Three of the most common phyla, Actinobacteria, Bacteroidetes, and Firmicutes all gained >20% deviance explained between model P+G and model P+G+H (Actinobacteria = 27.1%; Bacteroidetes = 24.6%, and Firmicutes = 25.2%; **Fig. 4B**; **table S6**). The most idiosyncratic features (i.e., those that gained >30% deviance explained by adding host-level factors), were microbiome PC2, the phylum Euryarchaeota, and the families Campylobacteraceae, Methanomethylophilaceae and Desulfovibrionaceae (**Fig. 4B**; **table S6**). Notably, even the most synchronous feature, microbiome PC1 (14% deviance explained by the P model), gained 23.2% deviance explained when adding host-level factors to the P+G model.

Removing covariates from model P+G+H one at a time, while keeping all other covariates intact, revealed that host identity explained nearly all of the deviance in our models (**Fig. 4C**; **table S6**; average loss in deviance explained by removing host identity = 17.3% compared to 0.2% deviance for all other factors). Beyond host identity, the next most important factor was the geographic area where the group traveled in the 30 days prior to sample collection, which on average, explained 1% of the deviance across all 52 features, with the strongest effects on microbiome PC1, Bifidobacteraceae, and Kiritimatiellaeota (**fig. S21**; **table S6**). The removal of all other individual predictor variables had only minor effects on deviance explained (**fig. S21**; **table S6**).

To investigate whether some of the idiosyncrasy we observed, especially at the host level, was due to host genetic effects, we tested for a relationship between the deviance explained by each GAM and the narrow-sense heritability (*h*^*2*^) of microbiome taxon abundance as estimated by Grieneisen et al.^33^. We found that higher levels of deviance explained by model P+G+H were predicted by higher taxon heritability (Pearson correlation: R=0.37, p=0.016; **Fig. 5A**). In contrast, we found no such effect at the population or group level, as expected since genotype is a property of individual hosts, not groups or populations (model P+G: R=0.047, p=0.76; model P: R=0.0085, p=0.96; **Fig. 5B**). In particular, we explained substantially more deviance by adding the host level to model P+G for microbiome taxa with *h*^*2*^ > 0.05 than we did for taxa with very low *h*^*2*^ values (model P+G+H: min=16.0, median=32.6, max=53.4 vs model P+G: min=4.6, median=11.1, max=26.8; **Fig. 5B**). These results suggest that some idiosyncrasy in gut microbiome dynamics is a consequence of differences in host genotype. However, because *h*^*2*^ estimates from the animal model cannot be mapped directly onto estimates of deviance explained in GAMs, direct estimates of genetic versus environmental effects on host dynamics remain an important topic for future work.

**Fig. 5.**
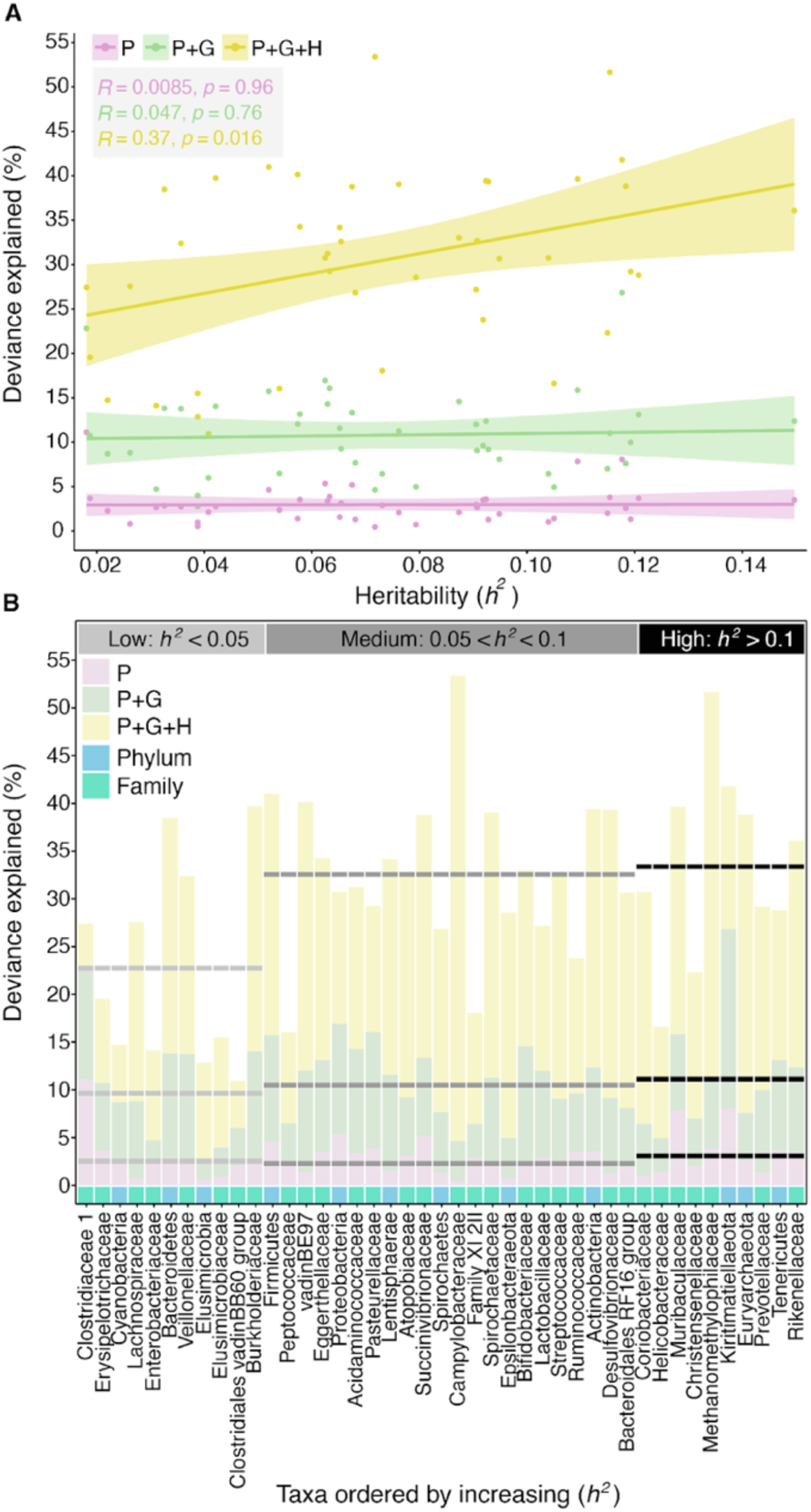
Microbiome taxon heritability is associated with idiosyncratic dynamics. (**A**) Deviance explained (y-axis) by the phylum and family level GAMs (from **Fig. 4**) plotted against the focal taxon’s heritability estimate (*h*^*2*^; x-axis). Pink, green and yellow denote model P, model P+G and model P+G+H, respectively. (**B**) Deviance explained (y-axis) across the model hierarchy (pink: model P; green: model P+G; yellow: model P+G+H) for each taxonomic feature (i.e., at the phylum and family level; x-axis). The x-axis is ordered by increasing heritability with light blue and turquoise squares representing phyla and families, respectively. Horizontal dashed lines show the average deviance explained per model for taxa with low heritability estimates (*h*^*2*^< 0.05; light gray); medium heritability estimates (0.05 < *h*^*2*^ < 0.1; dark gray); and high heritability estimates (*h*^*2*^ < 0.1; black).

### Gut microbial dynamics among social group members are more synchronized than for the host population at large

Previous research in humans and other social mammals, including the Amboseli baboons, finds that hosts in the same social group often have more similar gut microbiota than hosts in different social groups e.g.^1, 41, 58, 59, 62^. Likewise, in our current data set, several taxa exhibited abundances that were, on average, higher or lower within a given social group compared to their average abundance in the host population at large (**fig. S22, S23**). Hence, we tested whether shared social group membership is linked to greater microbiome community synchrony than hosts in different groups. In support, the patterns of temporal autocorrelation in **Fig. 3A** showed that hosts in the same group have detectably more similar microbiomes than those in different groups, especially for samples collected within 10 days of each other (**Fig. 3B**; Kruskal-Wallis: p < 2.2×10^−16^). Likewise, samples from the same group tended to occupy similar ordination space over time (**video S2**). While small, these group-level similarities were detectable, even for samples collected more than 2 years apart (**Fig. 3C**; **fig. S12A**). The addition of group-level splines to our GAMs led to gains in deviance that explained more than 10% for 15 of 52 microbiome features, including all three microbiome PCs, five phyla, and seven families (**Fig. 4B, 4C**; **table S6**). Several of these taxa were abundant in hosts, such as Firmicutes, Bacteroidetes, and Bifidobacteriaceae (**Fig. 4B, 4C**; **table S6**).

Because each social group has a somewhat distinctive gut microbiota, the effects of climate and diet on microbial dynamics may differ across groups. To test this idea, we added interaction effects between group identity and climate variables (rain and temperature), or between group identity and the first three PCs of diet to model P+G+H. However, these interactions did not lead to substantial gains in deviance explained in our models (**fig. S24**; **table S8**). For instance, adding the climate interactions explained on average an additional 0.95% deviance across all 52 features (range = −1.9% to 5.4%; **table S8**), and diet interactions explained, on average, an additional 1.2% deviance across all 52 features (range = −0.7% to 5.6%; **table S8**).

Gut microbial congruence among group members could also be linked to shared behaviors and environments: baboons in the same group eat the same foods at the same time, travel as a unit across the landscape, and may be grooming partners that are frequently in physical contact (**Fig. 1B, 1D**; **video S1**;^33, 36, 37, 38, 39, 40^). Indeed, after host identity, the next most important predictor variable in model P+G+H was the group’s home range in the 30 days before sample collection (**fig. S21**; **table S7**). Despite previous evidence for increased similarity in microbiome profiles among grooming partners in the Amboseli baboons^41^, we did not find evidence for this pattern in our current data set (**fig. S25**). Samples collected within 30 days of each other from individuals with strong grooming bonds were not substantially more similar than samples from animals with weak or no observed grooming relationship (mean Aitchison similarity between pairs with strong bonds = 0.645; mean Aitchison similarity between pairs weak or no bond = 0.646; **fig. S26**). Because of differences in methodology, the lack of a grooming effect in this data set should be interpreted with caution. Our prior research on this topic^41^ characterized microbial communities using shotgun metagenomic sequencing from >90% of social network members, all within 30 days of each other. In contrast, this current data set relies on 16S rRNA gene sequencing data from sparsely-sampled networks. Shotgun metagenomic data provide much higher taxonomic resolution than 16S rRNA identities, and may therefore more accurately capture the direct transmission between hosts.

## Conclusions

Using an unusually large time series dataset from a population of wild baboons, we found that gut microbial dynamics are both weakly synchronized across hosts and strongly idiosyncratic to individual hosts. Like members of a poorly coordinated microbial orchestra, microbial communities in different baboons are only weakly “in concert” across the host population. Instead, consistent with prior studies in humans and some wild animals e.g.,^1, 20, 21, 22, 23, 24, 25^, gut microbial dynamics are idiosyncratic at the level of individual hosts, and each baboon “player” approaches the gut microbial “song” differently. Our results contribute to mounting evidence that forces proposed to synchronize gut microbial metacommunities—shared environments, diets, and high rates of between-host microbial dispersal—can create modest synchrony among hosts, especially for hosts living in the same social unit. However, these forces are typically not strong enough to overwhelm powerful and well-known drivers of microbiome personalization, including host genetic effects, individual-level priority effects, horizontal gene transfer, and functional redundancy^16, 17, 18, 19^. Interestingly, these idiosyncratic dynamics were strong even for microbial phyla and families, whose dynamics reflect multiple microbial functions and interactions that potentially buffer them against large fluctuations in abundance. We expect that the personalized dynamics we observed will be even stronger for finer taxonomic levels, especially bacterial species or strains that exhibit a high degree of functional variability across hosts. In support, idiosyncratic dynamics were strongest for uncommon phyla or families (**fig. S16**), which might exhibit greater functional variation across hosts than common, abundant taxa.

Understanding the degree to which hosts in the same social group or population exhibit shared versus idiosyncratic gut microbiome dynamics may be useful to researchers interested in predicting individual microbiome changes, linking microbiome dynamics to health outcomes, and designing broadly effective microbiome interventions. These objectives have already been difficult to achieve, in part because of gut microbial personalization in humans and animals. For instance, predictive models of gut microbiome dynamics from one person have been shown to fail when they are applied to other people^27^. Although our focus here is on a single population of baboons, limiting extrapolation to other species and populations, our results provide important first-line evidence that microbiome predictions and interventions focused on microbiome taxa may require approaches that are either personalized or focus on microbial function, as opposed to taxonomic identities. Even then, “universal” microbiome therapies that work the same way for all hosts may be unattainable. Instead, microbiome interventions will likely work best when they are designed for host groups or populations that have shared compositions and dynamics. Functional redundancy and horizontal gene flow may also mean that functions will be more predictable than taxa, and prediction and intervention efforts that focus on microbiome functional traits (e.g., metabolite levels; the presence of specific functional pathways) will likely be less affected by gut microbiome personalization. Together, our results provide novel insights about the extent and ecological causes of microbiome personalization, and they indicate that personalized compositions and dynamics are not an artifact of modern human lifestyles and environments.

## Acknowledgments

We thank Jeanne Altmann for her essential role in stewarding the Amboseli Baboon Project, and in collecting and maintaining the fecal samples used in this manuscript. We also especially thank the members of the Maasai pastoralist communities in the Amboseli-Longido areas, on whose land we have lived and worked for 50 years. We also thank the Kenya Wildlife Service, Kenya’s Wildlife Research & Training Institute, the National Council for Science, Technology, and Innovation, and the National Environment Management Authority for permission to conduct research and collect biological samples in Kenya. We also thank the University of Nairobi, Institute of Primate Research, National Museums of Kenya, the Enduimet Wildlife Management Area, Ker & Downey Safaris, Air Kenya, and Safarilink for their cooperation and assistance in the field. We thank Karl Pinc for managing and designing the database. We also thank Tawni Voyles, Anne Dumaine, Yingying Zhang, Meghana Rao, Tauras Vilgalys, Amanda Lea, Noah Snyder-Mackler, Paul Durst, Jay Zussman, Garrett Chavez, and Reena Debray for contributing to fecal sample processing. Complete acknowledgments for the ABRP can be found online at https://amboselibaboons.nd.edu/acknowledgements/.

## Author contributions

EAA, JRB, LBB, RB, JAG, SM, and JT designed the research; EAA, SCA, RB, MRD, LG, JG, LRG, NG, SM, VY, NHL, TLW, RSM, JKW, LS, LBB, and JT, produced the data; JRB, TJG, DAWAMJ, LG, JCG performed the bioinformatics; JRB, KR, SM, performed the statistical analyses. EAA and JRB wrote the manuscript with important contributions from all authors.

## Data and materials availability

16S rRNA gene sequences are deposited on EBI-ENA (project ERP119849) and Qiita [study 12949, ^63^]. Analyzed data and code is available on the first author’s Open Science Framework / GitHub repository; for peer-review purposes, this is an anonymized link: https://osf.io/erdxa/?view_only=3323f05a5a9b479bac1124a5b07a62a9.

