## Supplementary Material for "Synchrony and idiosyncrasy in the gut microbiome of wild baboons"

#### Table of Contents

**1. SUPPLEMENTARY MATERIALS AND METHODS 2**

A. STUDY POPULATION 2
*Collection of baboon behavioral, ecological, and demographic data 2*
B. MICROBIOME DATA GENERATION 4
*Fecal sample collection and preservation 4*
*DNA extraction 4*
*PCR amplification, library construction, and Illumina sequencing 5*
*Analysis pipeline 5*
C. STATISTICAL ANALYSES 6
*Transformations to address compositionality 6*
*Principal Components Analysis 7*
*PERMANOVA on Aitchison distances 7*
*Linear mixed models 8*
*Autocorrelation analyses 8*
*Neutral model analysis 9*
*Generalized Additive Models 10*
Model P (population level) 11
Model P+G (Population + social group level) 11
Model P+G+H (Population + social group + host level) 12
GAM diagnostics and fit 12
Variable importance analysis 13
The effect of model complexity on deviance explained 13
*Taxon log-ratio analysis 14*

**2. SUPPLEMENTARY RESULTS 16**

A. SUPPLEMENTARY FIGURES 16
B. SUPPLEMENTARY VIDEO LEGENDS 42

**3. REFERENCES 43**

### 1. Supplementary Materials and Methods

#### A. Study population

Since 1971, the Amboseli Baboon Research Project (ABRP) has been collecting continuous, individual-based data on the members of several baboon social groups living in the Amboseli ecosystem in Kenya [1]. The population consists of primarily yellow baboons (*Papio cynocephalus*) with some admixture from neighboring anubis baboon (*P. anubis*) populations [note prior research finds no link between hybrid ancestry and the microbiome [2]]. The baboons studied by ABRP are individually recognized on sight, and full-time, experienced observers collect several types of data by visiting each group several times per week for half-day monitoring sessions (see the ABRP monitoring guide at https://amboselibaboons.nd.edu/downloads/).

##### Collection of baboon behavioral, ecological, and demographic data

We modeled how several variables predict microbiome composition and dynamics. At the host population-level, we modeled temporal changes in rainfall and temperature; at the social group-level we modeled group-specific data on home range use, diet, and group size; and at the host-level we modeled sex, age, social dominance rank, and grooming relationships. Below we explain how each data type was collected.

*Daily rainfall* and *daily maximum temperature* were collected using a rain gauge and min-max thermometer located in the ABRP field camp.

*Social group membership* and *group size* were known from census data collected during each monitoring visit to each social group. Census data record the identity of all study group members (infants, juveniles, adult females, and adult males) present in the group on the day of sampling. The study subjects were members of five primary social groups (Fig. 1).

*Group home range use* for each social group is known from GPS readings collected at the start of each half-day monitoring session and every 30 minutes thereafter until the observers departed the group.

*Group-level diet composition*. Data on diet composition were collected during 10-minute random order focal animal samples of adult female and juvenile behavior (*3*). Specifically, during each focal animal sample, we record activity (e.g. feeding, walking, resting) at 1-minute intervals; when feeding is observed, the food type is recorded. Food items are classified by species and part (e.g. *Ramphicarpa montana* blossoms). Food types were divided into 14 categories: (1) bark from all tree species (2) corms of all grass species; (3) pods from *Acacia tortilis* and *A. xanthophloea* (4) gum from *A. xanthophloea*; (5) seeds from *Acacia* trees; (6) blossoms from trees and shrubs, primarily from Acacia trees; (7) seed heads of all grass species; (8) *Azima tetracantha* fruit; (9) *Trianthema ceratosepala* fruit; (10) *Tribulus terrestris* fruit; (11) other fruits, including those from *Salvadora persica* and *Solanum dubium;* (12) non-grass leaves from e.g. *Acacia* trees, *Lyceum* sp., *Trianthema ceratosepala, Salvadora persica, Azima* *tetracantha*, *Rhamphicarpa montana*, and *Tribulus terrestris*; (13) non-corm components of all grass species; and (14) unknown/unidentified diet items (9% of all diet categories).

Following (*4, 5*), diet was measured using feeding observations on all adult females in the subject's social group in the 30 days prior to the sample collection date. We used these data to compute each social group's diet composition using (i) 30-day sliding windows with step size=1 day (Fig. 1D), and (ii) a PCA of clr-transformed group-specific diet composition (table S7). For the sliding windows, we first 'filled in' missing dates (i.e. dates with no diet recordings) with NAs to make the data time-continuous. Using a step size of 1, we then computed the diet composition for each social group in 30-day sliding windows. For the PCA, diet composition was computed for each social group and fecal collection date combination. More specifically, we computed the average relative abundance of each diet category from 1,000 random subsamples of all foraging observations recorded one month prior to the collection date. We then performed a PCA on clr-transformed diet compositions, and included all 13 principal components as predictor variables in the GAMs.

We used group-level diet data because our focal sampling data are too sparse to infer individual variation in diet over time-scales shorter than several months. Group-level diets are likely a sufficient representation of individual-level diets in the Amboseli population for three reasons (*5*). First, each baboon group travels together throughout the day, stopping to forage together in the same location and hence encountering the same resources at the same time. In support, previous analyses in Amboseli have found that group membership predicts diet composition, while dominance rank does not (*6*). Second, although individual baboons have dietary preferences (*7*), available food choices are determined by season and rainfall. For example, time spent foraging is more strongly predicted by environmental conditions than social group membership (*8*). Third, several studies that have intensively sampled individual-level diets over short time scales have found differences in the quantity but not composition of food consumed (*9, 10*).

*Host age* is known for any animal born into monitored study groups within a few days' error. For animals that immigrated into study groups from other social groups (~50% of adult males), age is estimated to within 1 to 3 years' error by comparing physical traits to known-aged males.

*Social dominance rank* for each baboon is known for each individual baboon in each month by assigning wins and losses in dyadic agonistic interactions between animals that live in the same social group. Wins are assigned when a focal baboon's opponent responded to the focal individual's aggressive or neutral gestures by giving only submissive gestures (in these cases, the opponent is also assigned a loss). These interactions are entered into a matrix of all same-sex individuals present for that month, in which individuals are ordered to minimize entries (counts of wins) below the diagonal. This procedure minimizes the cases in which a lower-ranking animal wins an encounter with a higher-ranking animal.

For a subset of our analyses, we also explored the predictive power of *grooming relationships*, which are recorded between individual baboons using representative interaction sampling during all group monitoring visits. During representative interaction sampling, observers move through the group as they perform focal animal samples on a randomized rotation of animals. The observer remains with each subject for 10 minutes and records all observations of physical contact for all individuals within their line of sight (including grooming interactions observed for animals who are not the subject of the focal sample). After 10 minutes, the observer moves to a new subject, ensuring representative sampling of the whole social group. For each grooming interaction, the observer records the identity of the groomer and the individual being groomed. We used these grooming interaction data to calculate a network of grooming bond strength, a dyadic grooming metric following [3]. Specifically, we calculated each individual's grooming relationships in the year prior to sample collection. Observed grooming bond strengths within each network were scaled to the highest number of interactions within each network such that the highest value in each group-year was 1 and the absence of a bond was represented by a zero. Hence, all grooming bonds were represented as a proportional strength relative to the strongest bond in their group in the hydrological year of interest.

#### B. Microbiome data generation

##### Fecal sample collection and preservation

Fecal samples were collected opportunistically from known baboons. Samples were typically collected within 15 minutes of defecation and were preserved in 95% ethanol in the field. Samples were stored at approximately 25°C until transport to the University of Nairobi, where they were freeze-dried, sifted to remove larger pieces of vegetation, and stored at -20°C. Upon arrival in the United States, samples were stored at -20°C. Samples were freeze dried because they were originally collected to measure steroid hormone levels e.g. [4, 5, 6, 7]. Previous work has shown that microbiome samples stored in ethanol and freeze-dried yield comparable results to other preservation methods [8].

##### DNA extraction

DNA was extracted using the MoBio and QIAGEN PowerSoil kit for 96-well plates. Prior to extraction, we aliquoted approximately 0.05 g of fecal powder from each of the 17,265 samples into 96-well plates. To minimize contamination during sample loading, we replaced the original sealing mats from the MoBio kit with Biotix silicone sealing mats. To load a given sample, we pierced the mat with a sterile pipette tip, inserted a polypropylene funnel into the well, and loaded ~0.05 g of fecal powder into e well. After all samples were loaded, the plate was sealed and stored at -80 °C until DNA extraction.

The resulting sample-loaded 96-well plates were extracted by the same person (MRD) at Argonne National Laboratory using the manufacturers' protocols, with a few modifications: (1) to hydrate the freeze dried samples and to minimize the risk of plate-clogging in the later stages of the extraction protocol, we increased the amount of PowerBead solution to 950 µL/well; (2) after the addition of the PowerBead solution and lysis buffer C1, we incubated the plates at 60°C for 10 minutes.

**PCR amplification, library construction, and Illumina sequencing**

Methods to construct 16S rRNA amplicon libraries were based on [9, 10]. We used polymerase chain reaction (PCR) to amplify an approximately 390 bp segment of the V4 region of the 16S rRNA gene using primers 515F - 806R, as follows:

**Forward**: GTGYCAGCMGCCGCGGTAA

**Reverse**: GGACTACNVGGGTWTCTAAT

Amplicons were quantified via Quant-iT PicoGreen dsDNA Assay Kit (ThermoFisher/Invitrogen cat. no. P11496). Equal amounts of amplicon DNA from each sample (70 ng) were pooled and cleaned using AMPure XP beads (Beckman Coulter). Typically, pools contained 1,623 samples from seventeen 96-well DNA extraction plates, but a few pools contained samples from sixteen plates. The cleaned amplicon pool was quantified using a Qubit fluorometer.

Libraries were sequenced on the Illumina HiSeq 2500 at the CHU Sainte-Justine Sequencing Core. We used the Rapid Run mode (2 lanes/run). Sequences were single indexed on the forward primer and 12bp Golay barcoded [which enables a high level of multiplexing [10]]. We sequenced 13 lanes split in two sequencing efforts: 8 lanes (on 4 flowcells) containing 12,912 samples were sequenced in March 2018 and another 5 lanes (on 3 additional flowcells) containing 7,776 samples in June 2018.

**Analysis pipeline**

To demultiplex the resulting sequences, we used bcl2fastq with the optional argument of −create-fastq−for-index-reads, which generates additional fastq files for index reads. The sample read masks are specified from the -use-bases-mask Y251,I12n*,Y251 option which specifies how each cycle is run. More specifically, the first Y251 means that we used the first 251 bases from the forward read; I12n* specifies that we used the next 12 bases for the index read and subsequently ignored the next n* bases. Finally, the last Y251 means that we used the first 251 bases from the paired reverse read. Following [11], index fastq reads were used for additional filtering of the samples to prevent barcode mis-assignment. We used DADA2's function filterAndTrim() to remove sequences in the index fastq files with a maximum error estimate, maxEE>0.1. Sequences in the sample fastq files were only retained if their corresponding index read passed the quality filter. Finally, Cutadapt [12] with minimum length=150 bases was used for Illumina library adapter removal and to remove reads that were too short for further analysis.

We used DADA2 [13] for sequence quality processing following the default protocol for large data sets (see https://benjjneb.github.io/dada2/bigdata.html) with the following exceptions in the FilterAndTrim() function: truncLen=c(240,160); maxN=0; maxEE=2; rm.phix=T; and truncQ=11. The FilterAndTrim()function filters the forward and reverse reads jointly, outputting only those pairs of reads that both pass the filter. This function call accomplished five tasks: (1) we truncated the forward and reverse reads at truncLen=c(240,160) nucleotides respectively; (2) we filtered out all reads with more than maxN=0 ambiguous nucleotides; (3) we filtered out all reads with more than maxEE=2 expected errors; (4) we discarded reads, rm.phix=T, that matched against the phiX genome; (5) we truncated reads at the first instance of a quality score less than or equal to truncQ=11. We used pooling (pool=T) to resolve exact ASVs (Amplicon Sequence Variants). This flag allows information to be shared across samples, which makes it easier to resolve rare variants that were present as singletons or doubletons in one sample but were present many times across samples. The DADA2 algorithm uses a parametric model of the errors introduced by PCR amplification and sequencing. Those error parameters typically vary between sequencing runs and PCR protocols. The LearnErrors()function provides a way to estimate those parameters from the data itself. DADA2 processing was performed on each HiSeq sequencing run of four flowcells with two lanes each (each lane equals a pool of libraries), and then combined with the function mergeSequenceTables(). Putative chimeric reads were removed with the function removeBimeraDenovo(). We used the Silva reference database, SILVA_SSU_r132_March2018.RData, for taxonomy identification using the IdTaxa()function from the R package DECIPHER [14]. We removed any ASVs that were assigned to chloroplasts, mitochondria, or eukaryotes.

For our final filtering steps, we included samples that met the following criteria: (i) the sample had a DNA extraction concentration that was at least 4X the blank on that sample's DNA extraction plate; and (ii) the sample returned more than 1,000 sequencing reads. Our final data set further filtered to ASVs seen at least 3 times in at least 20% of the samples.

#### C. Statistical analyses

##### Transformations to address compositionality

To address the compositional nature of our data [15], we applied the centered log-ratio (clr) transformation to all samples our count matrix [16] using the R package compositions [17]. This transformation places the data in a log-ratio coordinate space known as the Aitchison geometry in which standard statistical methodology can be applied [15, 16, 18]. The denominator of a log-ratio serves as the reference frame such that changes in the microbial population of one taxon are always relative to a reference frame given by one or multiple other microbial population(s) [19]. In the case of the clr-transformation, the reference frame is given by the geometric mean of the logtransformed abundances in each sample, meaning that the read count of each feature is compared to the read count of the 'average' feature of the same sample.

More formally, following [16], the clr-transformation for a sample can be obtained as follows

$$x_{clr} = [log(\frac{x_1}{G(x)}), log(\frac{x_2}{G(x)}), \ldots, log(\frac{x_D}{G(x)})]$$

$$G(x) = \sqrt[D]{x_1 \cdot x_2, \ldots, \cdot x_D}$$

where $x = [x_1, x_2, x_3, \ldots, x_D]$ denotes a sample containing $D$ "counted" features (here, ASVs), and $G(x)$ is the geometric mean of the focal sample, and serves as the reference frame. As $G(x)$ cannot be determined on sparse data, we added a small constant (0.65) to all of the elements of the abundance matrix (Justin Silverman, personal communication). An important property of the clr-transformation is scale invariance, which means that we can treat two communities as equivalent if they have the same relative abundances, even if they have different total abundances (e.g. sequencing depth) [15]. An important consequence of this property is that count normalization is unnecessary as it only leads to a loss of information and precision [15, 20]. Furthermore, the Euclidean distance between clr-transformed samples—the Aitchison distance [21]—also satisfies sub-compositional coherence which guarantees that the conclusions of an analysis on common taxa does not change if we decide to include rarer ones, and permutation invariance, which guarantees that the distance between two samples only depend on the compositional difference between them [15, 22, 23, 24].

#### Principal Components Analysis

To construct independent axes of microbiome community variation used in several analyses (e.g. the three microbiome principal components), we performed a principal components analysis (PCA) on clr-transformed relative abundances. While PCoA on distance matrices, such as Bray-Curtis dissimilarities, are commonplace in microbiome studies, this approach is not appropriate given the compositional nature the data. Instead, recent work on Compositional Data Analysis (CoDa) have shown that PCA on clr-transformed read counts or relative abundances is the appropriate form of ordination (see e.g. [15, 22, 24]). Performing PCA on clr-transformed read counts or relative abundances produces a Euclidean distance called the Aitchison distance, and when the distance metric is Euclidean, PCoA is equivalent to PCA (see e.g. [23]). We performed the PCA on the clr-transformed read counts using R's prcomp() function and extracted the first three PCs.

#### PERMANOVA on Aitchison distances

To identify the variables that explained variance in community composition across all samples, we ran two PERMANOVA analyses on our Aitchison distances via the adonis2()

function implemented in the R package vegan [25]. The first PERMANOVA tested the effect of season; its results are presented in fig. S3. The second PERMANOVA included 29 different population, group and host-level factors that could influence the microbiome (results are in table S4).

**Linear mixed models**

To test which microbiome features changed in abundance across seasons, we performed a differential abundance analysis to test which gut microbiome features increased/decreased in the wet as compared to the dry season (results are in Fig. 2C). Specifically, we fitted a linear mixed model to each of the 393 microbiome features (3 PCs; 3 alpha diversity metrics; 12 phyla; 34 families; and 341 ASVs), modeling season as a fixed effect, and host identity and social group as random effects. These models were fitted with the lmer() function from the R package lmerTest [26]. P-values were FDR corrected across all 393 linear models using the Benjamini-Hochberg procedure as implemented in the R function p.adjust(..., method="BH", n=393). The models were specified as follows:

$$\text{microbiome feature}_i \sim N\left(\alpha_{j[i],k[i]} + \beta_1(\text{season}_{\text{Wet}}), \sigma^2\right)$$
$$\alpha_j \sim N\left(\mu_{\alpha_j}, \sigma^2_{\alpha_j}\right), \text{ for host j} = 1, \ldots, \text{J}$$
$$\alpha_k \sim N\left(\mu_{\alpha_k}, \sigma^2_{\alpha_k}\right), \text{ for social group k} = 1, \ldots, \text{K}$$

where $\text{microbiome feature}_\text{i}$ represents the focal feature and $\alpha_j$ and $\alpha_k$ correspond to the random intercept for host $j$ and social group $k$, respectively. $\beta_1\,(\text{season}_{\text{Wet}})$ represents the regression coefficient for the effect of microbiome feature $i$ in the wet season as compared to in the dry season. Prior to clr-transforming the data, we averaged reads from samples collected from the same host animal in the same social group collected on the same date. The result of this analysis is presented in fig. 2C and tables S2 and S3.

**Autocorrelation analyses**

To test how microbiome communities changed as a function of time between sample collection dates, we measured patterns of temporal autocorrelation between samples collected from the same host, as well as between samples collected from different hosts living in the same or different social groups (results are in Fig. 3). Autocorrelation measures the similarity of two time points in the same time series at different time lags. To make the Aitchison distance more comparable to other ecological distances and more suitable for autocorrelation analysis, we re-scaled it to be bound between 0 and 1 and converted it to a similarity. Both procedures were done as follows:

$$\text{Aitchison similarity} = \left( \frac{1}{1 + \left( \frac{\text{aitchison distance}}{\text{max(aitchison distance)}} \right)} \right)$$

The resulting Aitchison similarity is bound between 0 and 1, where 1 corresponds to samples with identical compositions. Using the Aitchison similarity matrix containing all samples, we computed the average similarity of all samples from *(i) the same host in the same social group*; *(ii) different hosts in the same social group*; and *(iii) different hosts in different social groups* as a function of either the number of days or months between samples. For *(i)*, we only kept hosts with at least 2 samples. This analysis was performed through custom-written R scripts utilizing the R packages tidyverse [27], multidplyr [28] and parallel [29]. The results of this analysis are presented in figs. 3A, 3B and 3C.

Significance between these three comparisons *(i-iii)* was inferred by comparing their 95% confidence intervals (CIs). For example, if we were interested in comparing hosts in categories (ii) and (iii) above, we would subtract the latter 95% CI from the former. When this difference does not overlap zero, we can say that the two comparisons are significantly different from each other. These results are presented in fig. S11.

**Neutral model analysis**

To test whether dispersal limitation could explain microbiome idiosyncrasy, we estimated metacommunity-wide microbial migration probabilities in each season and year using the Sloan Neutral Community Model for Prokaryotes [30, 31]. This model assumes that each host's microbial composition (i.e., local community) is the outcome of stochastic population dynamics and microbial immigration from other hosts in the microbiome metacommunity (i.e., other local communities). Briefly, local communities have a constant size $N$, and individual microbes within each local community die at a constant rate. These deaths create vacancies that can be occupied, either by individuals immigrating from the microbiome metacommunity (with probability $m$), or by the offspring from any taxon within the local community (i.e., from reproduction within the same host, with probability $1\text{-}m$). Species that are common in the metacommunity have a higher chance of occupying vacancies than rare species. Without immigration from the microbiome metacommunity, ecological drift leads each host's microbial diversity to reduce to a single taxon. Thus, the migration probability, $m$, represents the metacommunity-wide probability that any taxa, randomly lost by drift from local communities, will be replaced by dispersal from the microbiome metacommunity, as opposed to reproduction within hosts [30, 31]. Following Burns et al. [32], $m$ can be interpreted as a measure of dispersal limitation with low migration probabilities being equal to high dispersal limitation. We estimated metacommunity-wide values for $m$ in each season and hydrological year by treating individual hosts as local communities, and defining the microbiome metacommunity as either the hosts' social group or the whole host population. We fit the neutral models using nonlinear least-squares regression as implemented in the R package tyRa [33]. The results of the analysis are presented in fig. S16.

*Defining each social group as the metacommunity*. We started by averaging reads from samples collected from the same host animal in the same social group collected on the same date. We then rarefied the abundance matrix containing all samples. For each social group, we then selected one random sample per host animal, hydrological year, and season; for example, if there were 49 host animals from Nyayo's group with a total of 456 samples in the 2002's wet season, a subsetted abundance table containing 49 samples from 49 hosts was created. This formed the set of local communities for the focal social group and time period. All samples from the focal group and time period were used to create the metacommunity (continuing with the above example, it would contain 456 samples from 49 hosts). The neutral model was then fit using the tyRa::fit_sncm(spp=…, pool=…) function with the spp and pool arguments corresponding to the set of local communities and the metacommunity, respectively. This procedure was permuted 500 times. Thus, for each permutation one model was fit for a combination of social group, hydrological year and season. The parameter $m$ was extracted for each model. The mean and standard deviation was computed across all permutations.

*Defining the whole host population as the metacommunity*. We started by averaging reads from samples collected from the same host animal in the same social group collected on the same date. We then rarefied the abundance matrix containing all samples. For each host, we sampled one random sample from the whole host population per hydrological year and season; for example, if there were 2000 samples across the population from 200 hosts in the wet season of year 2002, a subsetted abundance table containing 200 samples from 200 hosts was created. This formed the set of local communities for the focal time period. All samples from the focal time period were used to create the metacommunity (continuing with the above example, it would contain 2000 samples from 200 hosts). The neutral model was then fit using the tyRa::fit_sncm(spp=…, pool=…) function with the spp and pool arguments corresponding to the set of local communities and the metacommunity, respectively. This procedure was permuted 500 times. Thus, for each permutation one model was fit for each combination of hydrological year and season. The parameter $m$ was extracted for each model. The mean and standard deviation was computed across all permutations.

#### Generalized Additive Models

Generalized Additive Models (GAMs) are a flexible family of models that do not assume a linear relationship between the response and predictor variables [34]. The impact of any predictor variable is captured through a so-called smooth function which, depending on the underlying patterns in the data, can be either linear or nonlinear [34]. To balance the overfitting/underfitting trade-off, GAMs allow both the smoothness (i.e., how wiggly a given function is) and the effect size of component functions to be controlled through regularization/penalization [34, 35].

The modeled data are hierarchical as microbiome samples can be grouped into those collected from individual hosts, which themselves can be further grouped into social groups, with variables measured for hosts, social groups and the population as a whole. Hence, we constructed three hierarchical models to describe temporal trends at the (i) population level (model P); (ii) population and the social group level (model P+G); and (iii) population, social group level, and the individual host level (model P+G+H). GAMs were fitted to a subset of the full data, consisting of 4,277 16S rRNA gene sequencing profiles from the best-sampled 56 baboons living in the 5 main social groups between 2002 and 2010 (fig. S2; min=48; median = 72.5; max = 164 samples). We fit GAMs using the R function bam() from the package mgcv which uses fast restricted maximum likelihood (method="fREML") as the smoothing parameter estimation method [34]. We fitted each GAM (i.e., model P; model P+G; and model P+G+H) to 52 different gut microbiome features describing different aspects of the gut microbiome: three principal components (PC1-PC3) and three alpha diversity indices (species richness, exponent of Shannon diversity, and the inverse Simpson index) to describe community-level changes, as well as clr-transformed relative abundances of 12 phyla and 34 families. The results of these models are presented in fig. 4A and table S6.

To minimize residual autocorrelation, we fitted the residuals of a Gaussian process to each GAM. This was done by first making each of the modeled host's time series (fig. S2) continuous by 'filling in' missing collection dates (i.e. days) with NAs, and then fitted a Gaussian process via the R function gausspr() from the package kernlab [36]. The input to the gausspr() function was one of the 52 gut microbiome features from a focal host. We then extracted and used the residuals from the Gaussian process as the response variable in each GAM.

All response variables are assumed to be distributed $N(u_i, \boldsymbol{\sigma}^2)$ where $E(Y_i)=u_i$ is the identity link, and index $i$ indexes the $i-th$ sample. The three models are defined as follows:

***Model P (population level)***

$$g(\mu_i) = \underbrace{f_1(\texttt{Month}_i) + f_2(\texttt{HydroYear}_i)}_{\text{Population level}} \qquad \text{(model P)}$$
$$i = 1, 2, \ldots, 4,277$$

where $f_{1-2}$ are smooth functions for the $i-th$ month and hydrological year, respectively. The population-level variables included in model P were monthly and annual total rainfall and average monthly and annual maximum temperature (table S5).

***Model P+G (Population + social group level)***

$$g(\mu_i) = \underbrace{f_1(\texttt{Month}_i) + f_2(\texttt{HydroYear}_i)}_{\text{Population level}} + \underbrace{f_3(\texttt{Time}_i, \texttt{Group}_{k[i]})}_{\text{Social group level}} \qquad \text{(model P + G)}$$
$$i = 1, 2, \ldots, 4,277; \quad k = 1, 2, \ldots, 5$$

where $f_{1-2}$ are smooth functions for the $i-th$ month and hydrological year, respectively. The new term $f_3$ specifies a social group-level smooth as an interaction between time (numerical representation of collection date) and social group with $k[i]$ indexing the $k-th$ group in which the $i-th$ sample was collected in. Beyond the population-level covariates from $\text{model P}$, here we also accounted for social group size, each social group's total home range area and fraction unique home range area, a 3-dimensional interaction between latitude, longitude and time, as well as 13 different diet PCs as potential drivers of the gut microbiota (table S5).

##### *Model P+G+H (Population + social group + host level)*

$$g(\mu_i) = \underbrace{f_1(\texttt{Month}_i) + f_2(\texttt{HydroYear}_i)}_{\text{Population level}} + \underbrace{f_3(\texttt{Time}_i, \texttt{Group}_{k[i]})}_{\text{Social group level}} + \underbrace{f_4(\texttt{Time}_i, \texttt{Host}_{r[i]})}_{\text{Host level}} \qquad (\text{model P + G + H})$$

$$i = 1, 2, \ldots, 4,277; \quad k = 1, \ldots, 5; \quad r = 1, \ldots, 56$$

where $f_{1-2}$ are smooth functions for the $i-th$ month and hydrological year, respectively, and $f_3$ specifies a social group-level smooth as an interaction between time (numerical representation of collection date) and social group with $k[i]$ indexing the $k-th$ group in which the $i-th$ sample was collected. The new term $f_4$ specifies a host-level smooth as an interaction between time and host with $r[i]$ indexing the $r-th$ host from which the $i-th$ sample was collected. Finally, in addition to the covariates that were included in $\text{model P + G}$, here we also accounted for each host's sex, age and social dominance rank (table S5).

##### *GAM diagnostics and fit*

We evaluated model fit focusing on two diagnostic plots: (1) QQ-plots of residuals, and (2) deviance residuals vs. fitted values. For some responses, at low values, the deviance residuals were larger than predicted by the theoretical quantiles. In most cases, there were no patterns in the residual vs. fitted values, apart from some clusters of outliers in some of the plots. Overall, the majority of models and response variables produced reasonably good fits to the data (table S10). We also focused on model diagnostics from the mgcv::k.check() function, which tests whether the basis dimension for each smooth in the focal model is adequate (table S10). Regardless of the response variable, we always used the same number of basis dimensions (i.e., *k*, the maximum allowed degrees of freedom) for each smooth term. Moreover, for model P+G+H and model P+G, the number of basis dimensions for the smooth term(s) defining the previous hierarchical level(s) always remained the same. For model P, we always used a cyclic intra-annual smooth (bs="cc") with *k*=7 and an s-smooth for hydrological year with *k*=8. To strike a balance between the degrees of freedom, computation, and generality across response variables (i.e., goodness-of-fit across response variables), *k* was set to 20 and 60 for the s(time, by=group) and s(time, by=host) smooths in model P+G and model P+G+H, respectively. This allowed us to compare fits (e.g., deviance explained) across hierarchical levels and response variables. However, as a consequence, each model was not necessarily 'optimized' for the focal response variable. As seen in table S10, while the *k-index* is below one, the maximum degrees of freedom (*k'*) is always lower than the effective degrees of freedom (*edf*) which indicates that the basis dimension for each smooth term was sufficiently large.

While not directly related to the choice of basis dimensions, the *m* parameter specifies the order of the penalty for a smooth term: *m*=2 is the default for many basis functions, but for hierarchical GAMs with nested global and group-level smoothers, it is recommended to set *m*=1 for the group-level smoother. This will penalize the squared first rather than the squared second derivative, and a group-level smoother with *m*=1 will have a more restricted null space, ultimately reducing collinearity with the global smoother (see [37] for more details). As we modeled three hierarchical levels, we set *m*=2 for the population and social group-level smoothers, and *m*=1 for the host-level smoother.

Lastly, to increase computational efficiency, we utilized (1) the bam() function which uses fast restricted maximum likelihood (method="fREML") as the smoothing parameter estimation method [34]; (2) discrete=TRUE, which discretizes the covariates (see [34, 38] and Wood and Li 2019 for detailed information); and (3) parallel computing using the argument nthreads(24,1), which means we used 24 threads for discrete inner products, and 1 for code calling BLAS (see ?mgcv::bam() for more details). We re-ran our P+G+H models with discrete =FALSE, and results were nearly identical.

##### *Variable importance analysis*

To quantify how much each independent variable (n=28) in model P+G+H contributed to deviance explained, we removed covariates from the model one at a time, while keeping all other covariates intact. We performed this procedure for each response variable (n=52), thus running a total of 1,456 models. This analysis only became feasible when arguments discrete=TRUE and nthreads(24,1) were used. The results from this analysis are presented in figs. 4C and S18, and table S8.

##### *The effect of model complexity on deviance explained*

In many settings, increasing the complexity (i.e., degrees of freedom) of a model also increases variance or deviance explained, even if additional model parameters do not improve model fit. To ensure that this explanation does not account for our observation that the hierarchical models explain more deviance than single level models (e.g., P+G versus P), we performed two analyses. These are explained in detail below, but briefly, in the first, we randomized either social group or host identity, including group and host-level covariates, and tested whether these randomized models led to a decline in deviance explained. In the second analysis, we performed a simulation experiment to investigate the effect of model complexity on deviance explained.

Randomization analyses. For each principal component (PC1, PC2, and PC3), we ran model P+G+H 10 times, each time randomizing social group membership and group-level variables (group size, home range variables, and diet PCs) across samples, while keeping each sample's population- and host-level variables intact. We extracted the deviance explained for each model and averaged across the 10 permutations. These results are presented in Fig. S18 ('Average of group label permutations'). We then ran model P+G+H 10 times, each time randomizing host identity and other host-level variables (sex, age, rank) across samples while keeping each sample's annual, seasonal, and group identity intact. We extracted the deviance explained for each model and averaged across the 10 permutations. These results are presented in Fig. S18 ('Average of host label permutations').

*Simulation*. We simulated data with a strong population-level effect, but with very similar average functional responses among social groups. We simulated data from 4 hosts, each with 100 samples divided into two social groups. The population-level effect, which was assigned to all 4 host individuals was drawn from a normal distribution with a mean equal to -1 and +1 in the first and last 50 time points, respectively. On top of this, group level noise which was drawn from a normal distribution with a zero mean and a standard deviation of 0.5 was added to the two hosts in each social group. This experiment aims to specifically test if explanatory power (i.e., deviance explained) increases when a model (such as model P+G+H) has several observations to reconcile (at any given time-point) at the group level but only one at the host level (which only contains noise). We fit three models mimicking model P, model P+G and model P+G+H using mgcv::bam(…, select=F, method="ML"). The results of this analysis are presented in Fig. S19.

**Taxon log-ratio analysis**

To understand how individual microbiome phyla and families contributed to host- or group-level idiosyncratic abundances, we constructed log-ratios comparing each taxon's clr-transformed relative abundance in a given host or social group to its clr-transformed relative abundance in the host population at large. These log-ratios were constructed by subtracting each focal taxon's average clr-transformed relative abundances across all samples belonging to a given host or social group from its grand mean across the host population (i.e., the pooled average of the average clr-transformed samples from each host; i.e. one average clr-transformed value per host). We also performed this analysis within each season and year, but for simplicity we show the cross-sectional analysis below. Prior to running these analyses, we averaged the relative abundance of samples collected from the same host in the same group collected on the same date. The results of this analysis are presented in figs. 3E and 5.

More formally, let $\widehat{x_{i,k}^{clr}}$ denote the average clr-transformed relative abundance for taxon $i$ in host or social group $k$, which is given by

$$\widehat{x_{i,j}^{clr}} = \frac{1}{n}\sum_{j=1}^{n} x_{k,j}^{clr} = \frac{x_{k,1}^{clr} + x_{k,2}^{clr} + \cdots + x_{k,n}^{clr}}{n}$$

Then, the log-ratio of taxon $i$'s average clr-transformed relative abundance in host or social group $k$ to its average clr-transformed relative abundance in the baboon population at large, is given as follows

$$log\left(\frac{x_{i,k}}{x_i}\right) = \frac{1}{n}\sum_{j=1}^{n} x_{k,j}^{clr} - \frac{1}{n}\sum_{j=1}^{n} x_j^{clr}$$

A positive/negative log fold change corresponds to taxa whose relative abundance was higher/lower in the focal host or social group compared to in the host population at large.

### 2. Supplementary Results

#### A. Supplementary figures

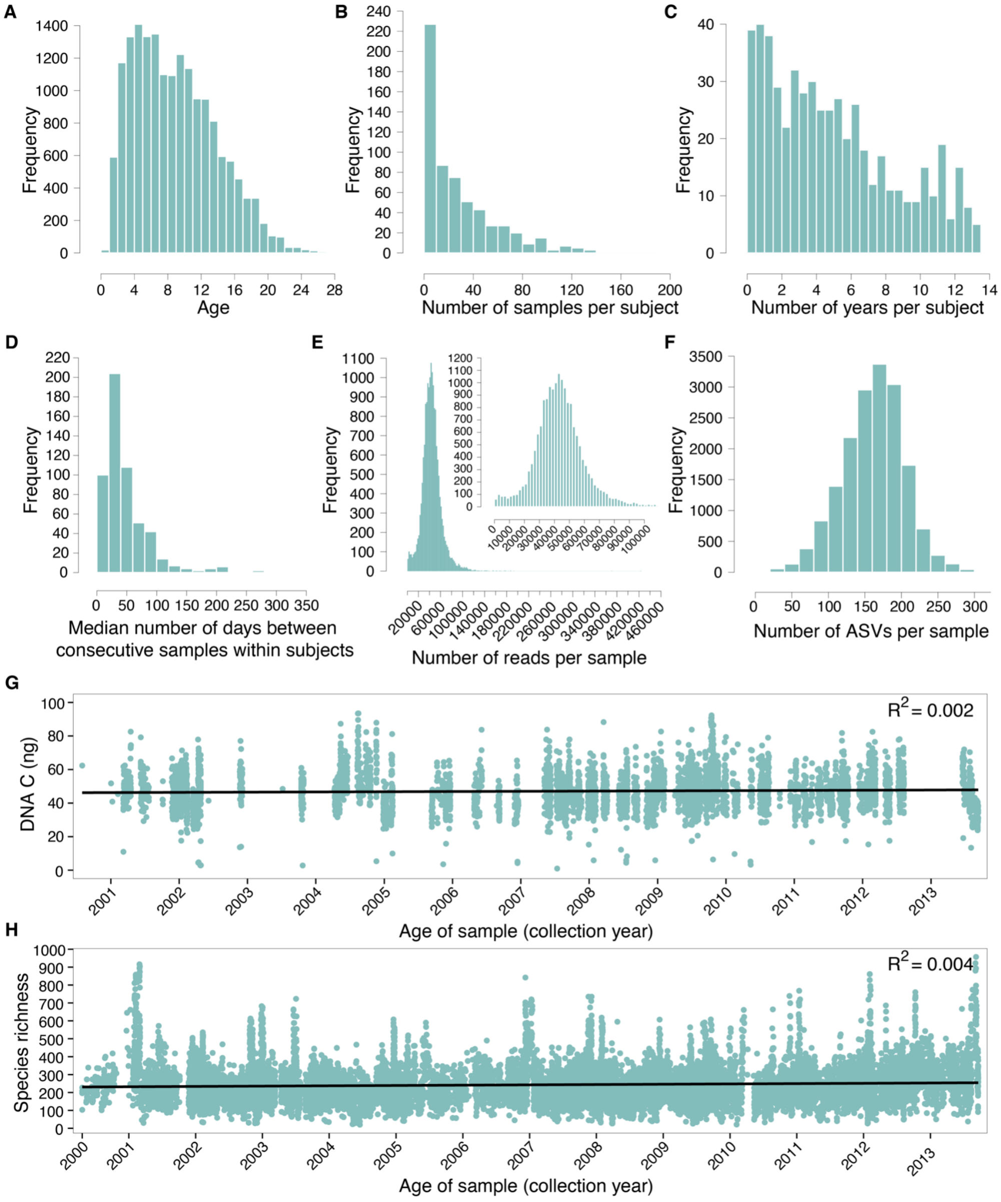


**Fig. S1.** Histograms showing sample distributions for all 17,265 16S rRNA gut microbiome profiles from 600 hosts. Plots show **(A)** the number of samples available for hosts of different ages; **(B)** the number of samples per subject; **(C)** the number of years of sampling per subject; **(D)** the median number of days between consecutive samples within subjects; **(E)** the number of sequencing reads per sample (the inset figure shows the histogram without the long tail); and **(F)** the number of ASVs per sample. Sample age does not predict **(G)** DNA concentration (G; $R^2$=0.002), or (H) species richness (richness was calculated prior to filtering; $R^2$=0.004), indicating that older samples were not more degraded.

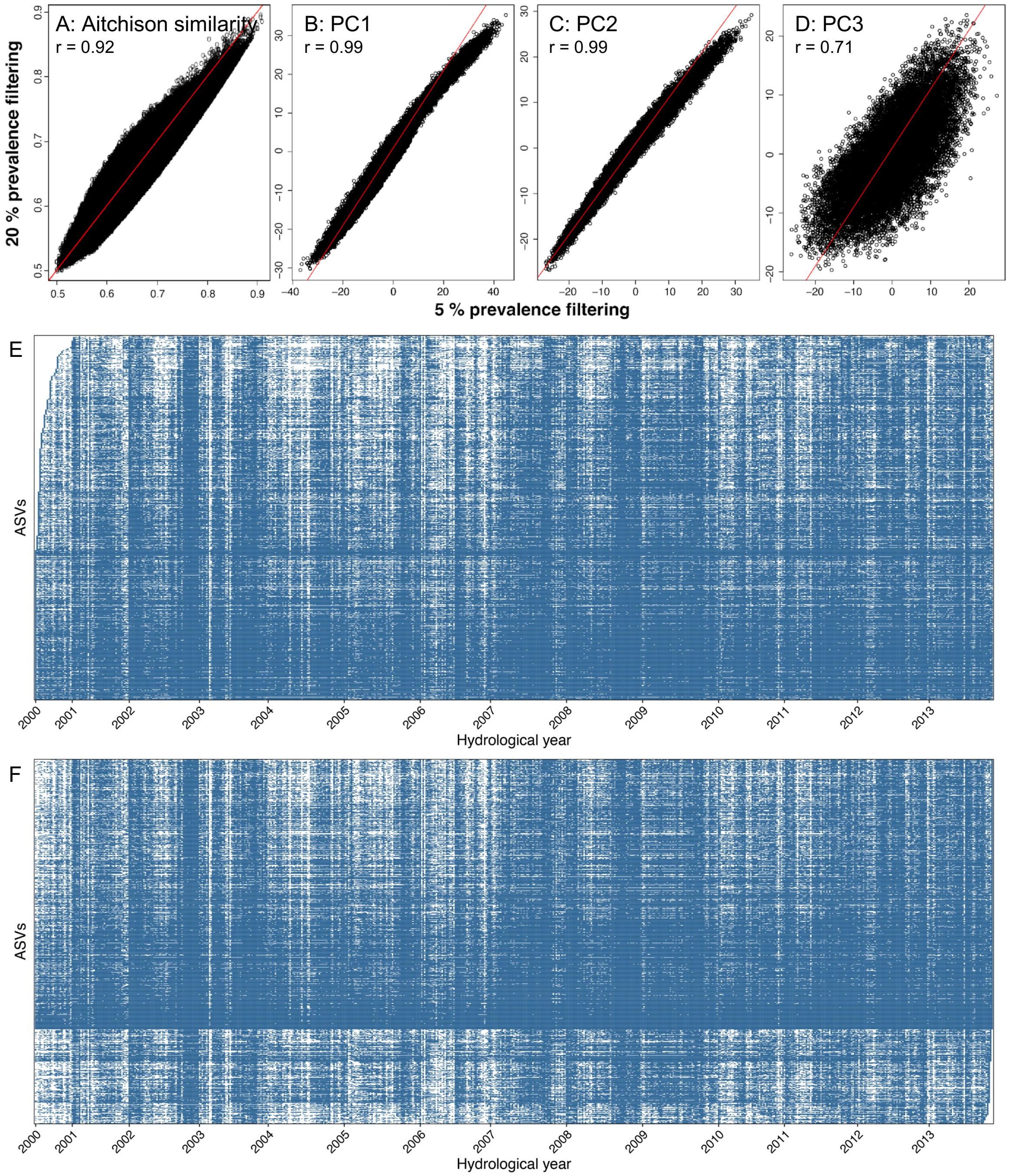


**Fig. S2** Comparison of community metrics for the microbiome data set filtered to 20% versus 5% prevalence across samples. **(A-D)** Correlations and Pearson's r values for Aitchison similarities, PC1, PC2, and PC3 for data sets filtered to 5% vs 20% prevalence across samples. **(E)** and **(F)** are heat maps to check for evidence of ASV extinction or recent invasion across the 14 years of the data set. Each row represents one of 805 ASVs present in at least 5% of samples. Each column represents one week of sampling between 2000 and 2013. A box is shaded blue if a given ASV was detected in any sample the population in that week; the box is white if the ASV was not detected in any sample in that week. The ASVs are sorted by **(E)** the first week or **(F)** the last week they appeared in any sample in the data set.

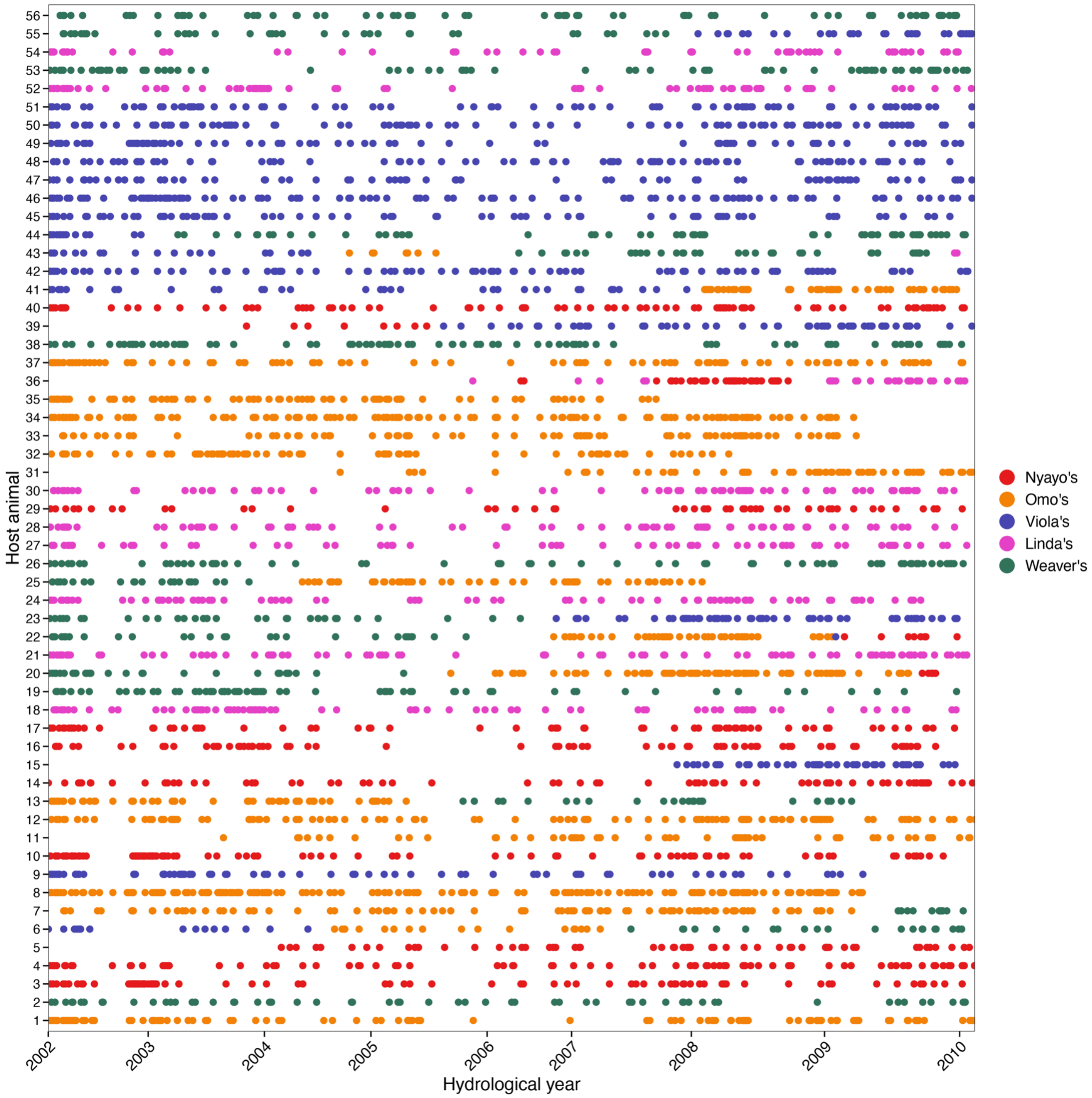


**Fig. S3.** Dot plot showing 4,277 samples from the best-sampled 56 baboons between 2002 and 2010. These samples are a sub-set of the data in fig. S1 and were used in our generalized additive models (GAMs; min=48; median = 72.5; max = 164 samples). Each row on the y-axis shows a unique host animal, and the x-axis shows the fecal sample collection date grouped by the hydrological year (November 1 to October 31) when the fecal sample was collected. Dot color corresponds to the 5 original social groups (see legend; colors are the same as in Fig. 1). Adult males disperse between social groups, and dispersals by males can be identified by cases in which colors in the same row (corresponding to the same unique individual) change over time (e.g. host animal 7, 25, and 39).

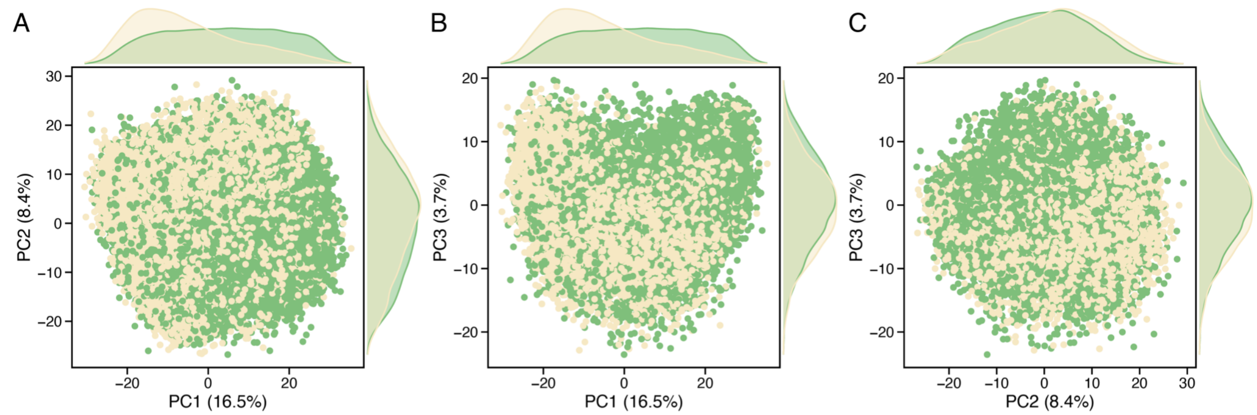


**Fig. S4.** Principal components analysis (PCA) of Aitchison distances between all 17,265 samples. Each dot is colored by the season it was collected in, wet (green) or dry (yellow). Density plots along the x- and y-axes depict sample distributions in each season. Each panel shows a combination of the first three PCAs: **(A)** PC1 and PC2; **(B)** PC1 and PC3; and **(C)** PC2 and PC3. A PERMANOVA showed that season was a significant factor explaining variation in the baboon gut microbiome ($p<0.001$; $R^2 = 0.0194$).

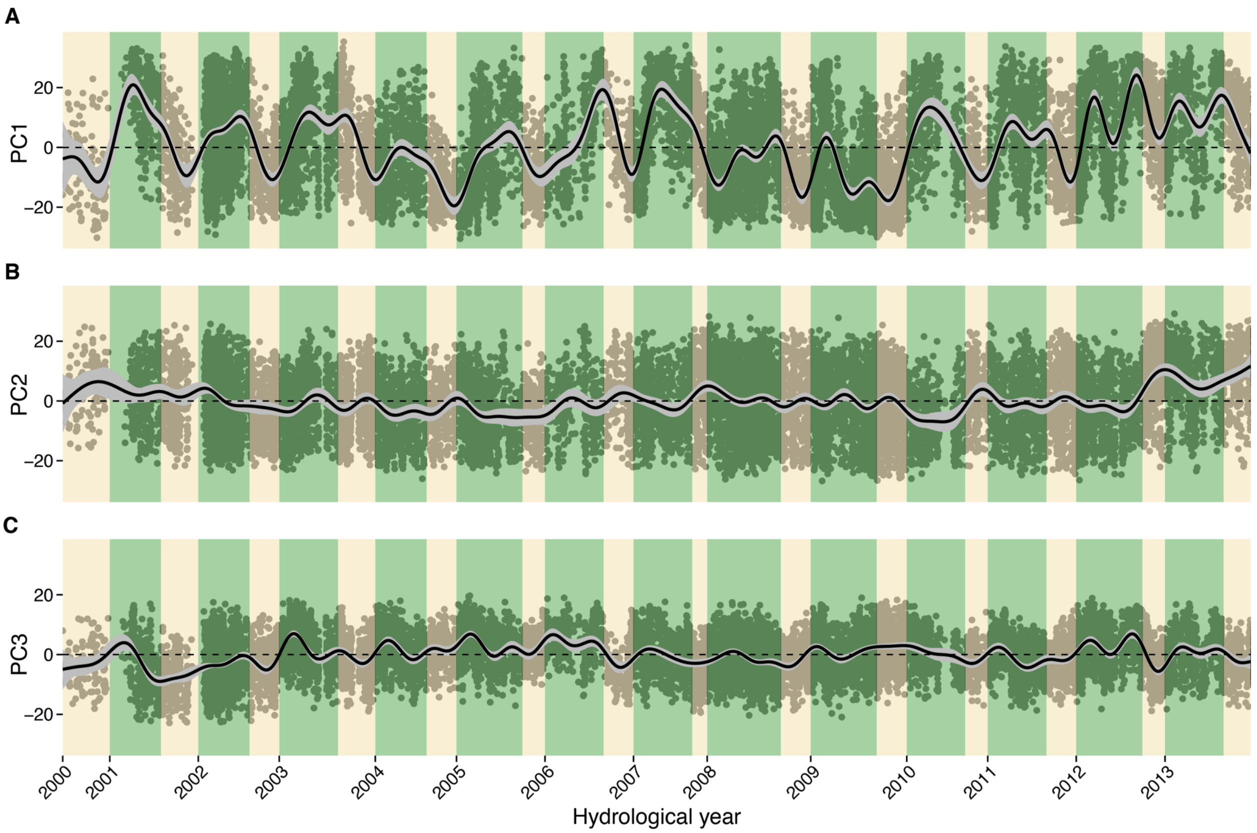


**Fig. S5.** The first three microbiome PCs for all 17,265 samples plotted over time (see fig. S3 for ordination plots of the same PCs). Panels **(A-C)** depict PC1-PC3 plotted over time; points show PC estimates for individual samples; the black line shows fitted population-level smooths in black; the grey bands show 95% confidence intervals. Each year on the x-axis reflects a hydrological year; green panels correspond to the wet season and yellow panels correspond to the dry season.

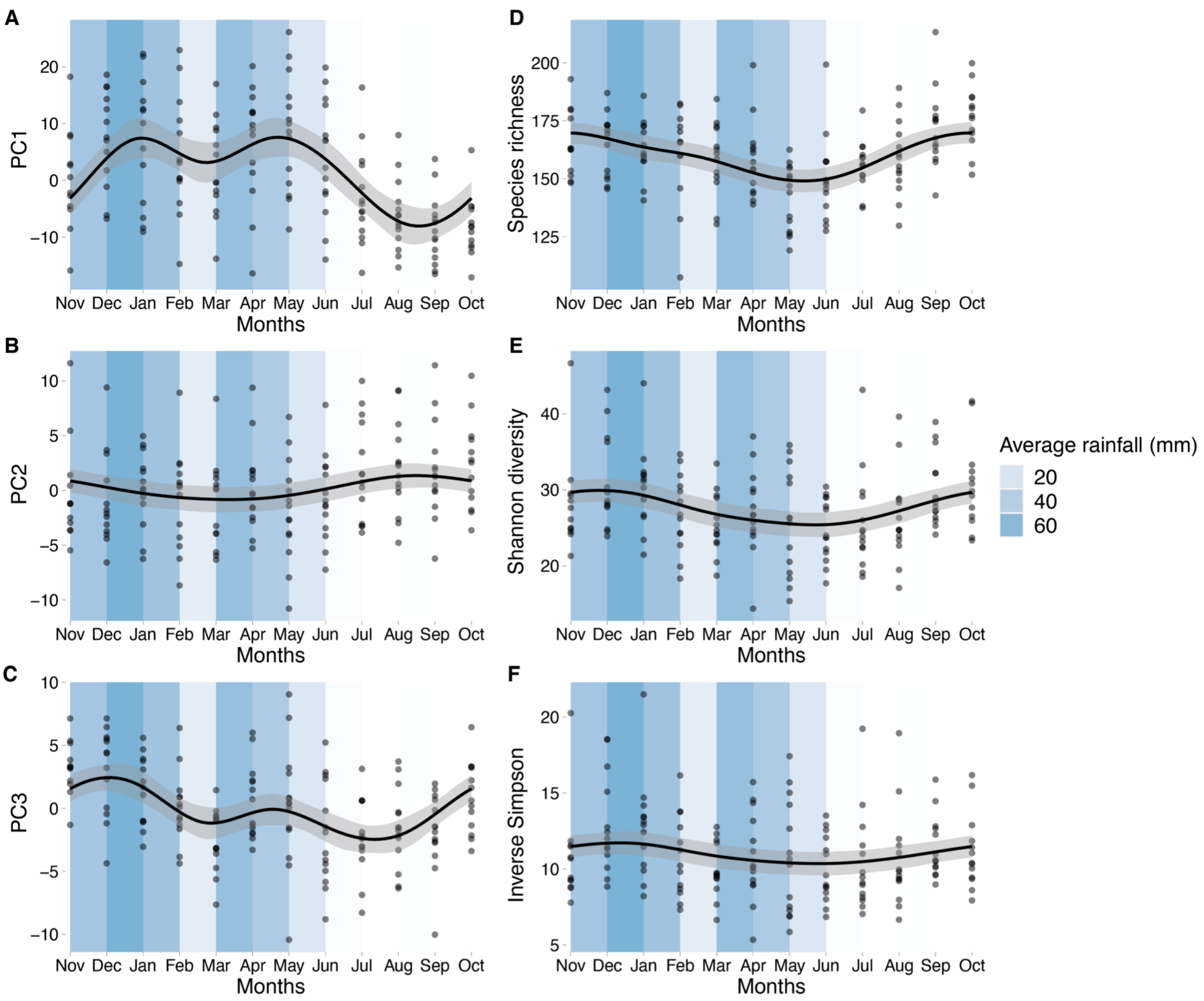


**Fig. S6.** Intra-annual changes in **(A-C)** the top three microbiome PCs, and **(D-F)** three measures of alpha diversity. Each dot represents the mean PC or alpha diversity measure for a given month and hydrological year (November to October). Black lines and grey bands depict fitted smooth and 95% confidence intervals. Blue panels on each plot represent the average rainfall in a given month across the 14 years of the data set.

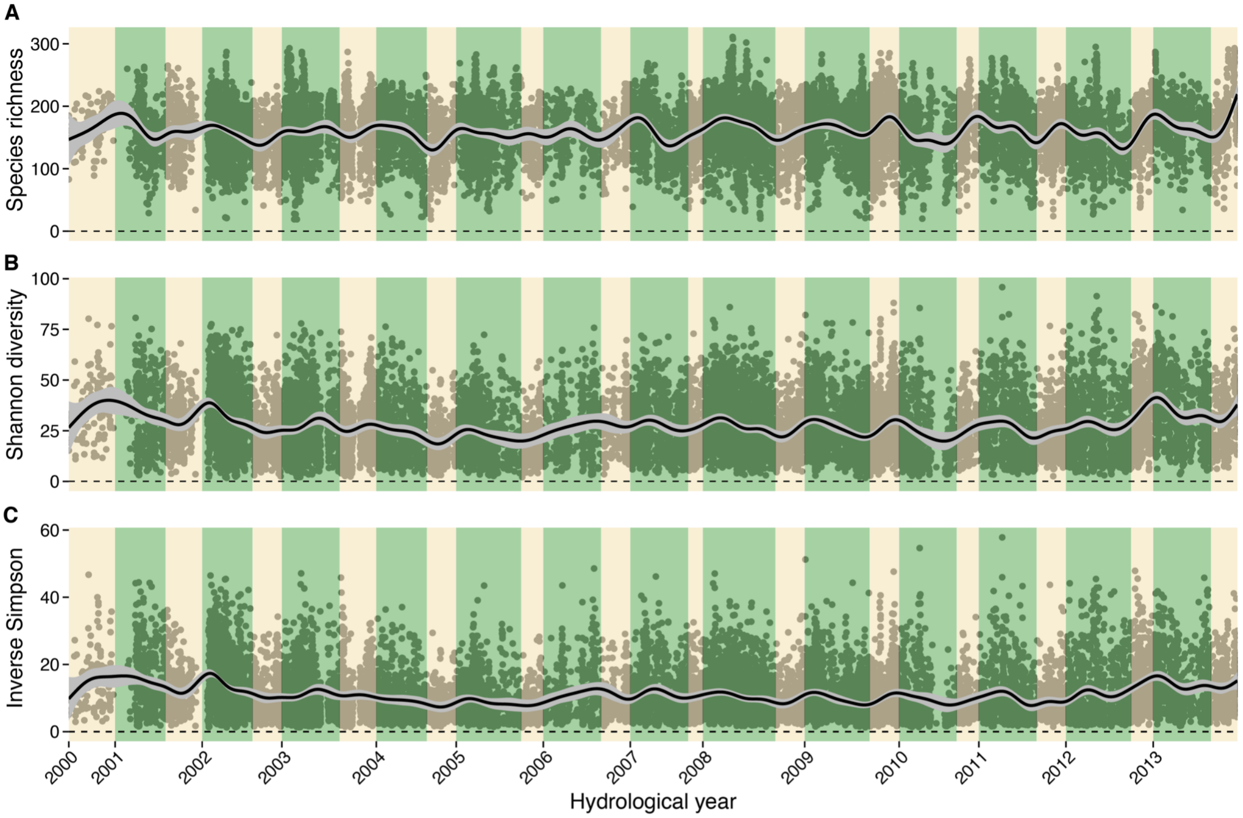


**Fig. S7.** Temporal dynamics of microbiome alpha diversity in all 17,265 samples. Alpha diversity was measured as: **(A)** Species richness (mean=162 taxa; sd=42.2; cv=0.26), **(B)** Shannon's diversity (mean=27.8 taxa; sd=14.3; cv=0.51), and **(C)** the Inverse Simpson index (mean=11 taxa; sd=7.3; cv=0.66). Each year on the x-axis reflects a hydrological year; green panels correspond to the wet season and yellow panels correspond to the dry season. Points show alpha diversity estimates for individual samples; black lines and grey bands show smooths and 95% confidence intervals.

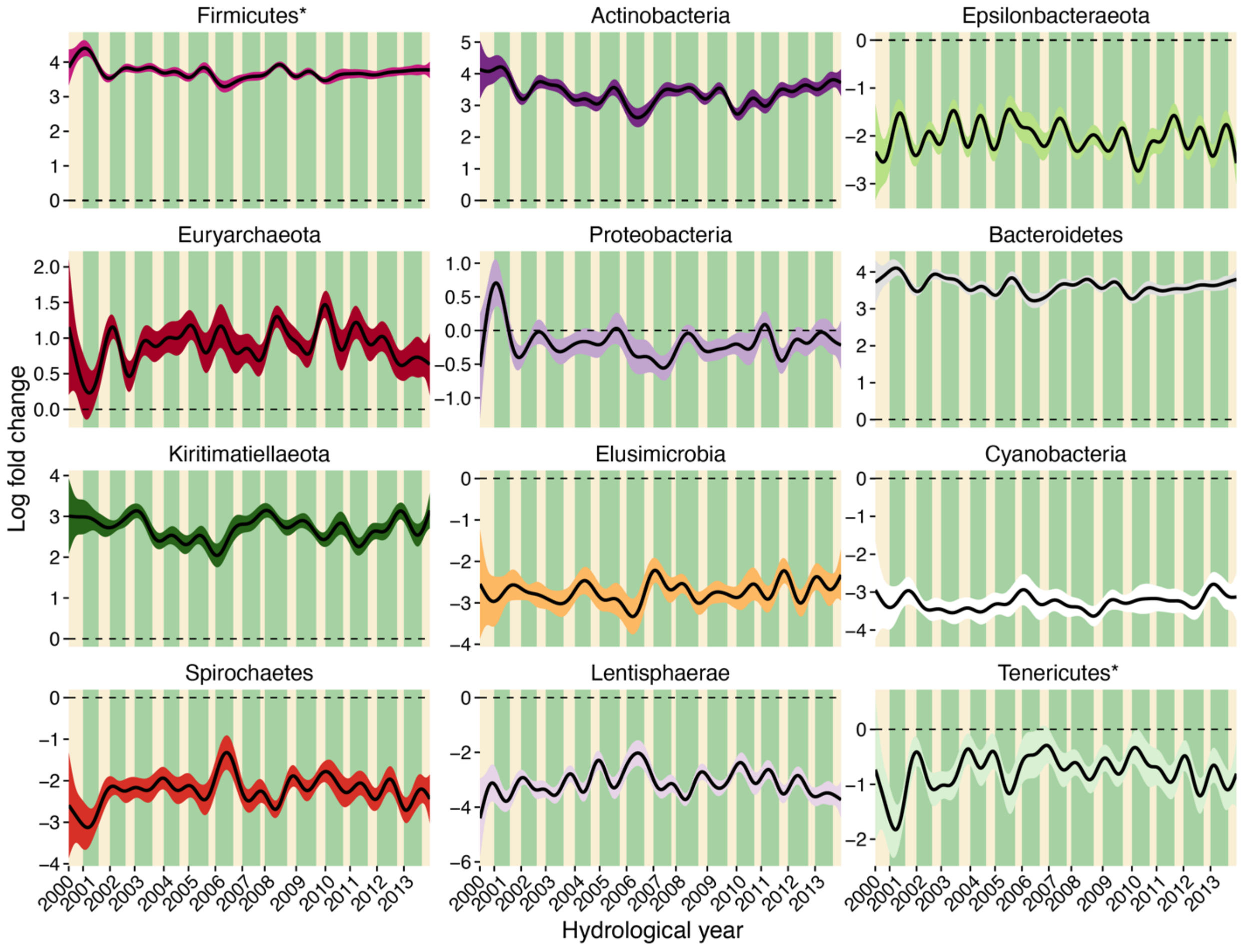


**Fig. S8.** Change in clr-transformed relative abundances across seasons and years for 12 bacterial phyla present in >20% of samples. Phyla are presented in the same order as in Fig. 2C, starting with those that have the largest log fold increase in the wet season and ending with those that have the largest log fold increase in the dry season. Changes in the clr-transformed relative abundances are expressed as log fold changes, which represent changes in the focal taxon's relative abundance compared to changes in the relative abundance of an average taxon in the same microbial community. A positive/negative log fold change corresponds to taxa whose relative abundance is higher/lower than the average taxon in the same microbial community. Black lines represent fitted smooths; colored bands represent 95% confidence intervals. Each year on the x-axis reflects a hydrological year; green panels correspond to the wet season and yellow panels correspond to the dry season. Asterisks denote taxa that were significant (FDR threshold = 0.05 for n = 393 models) in the linear models in Fig. 2C and table S2.

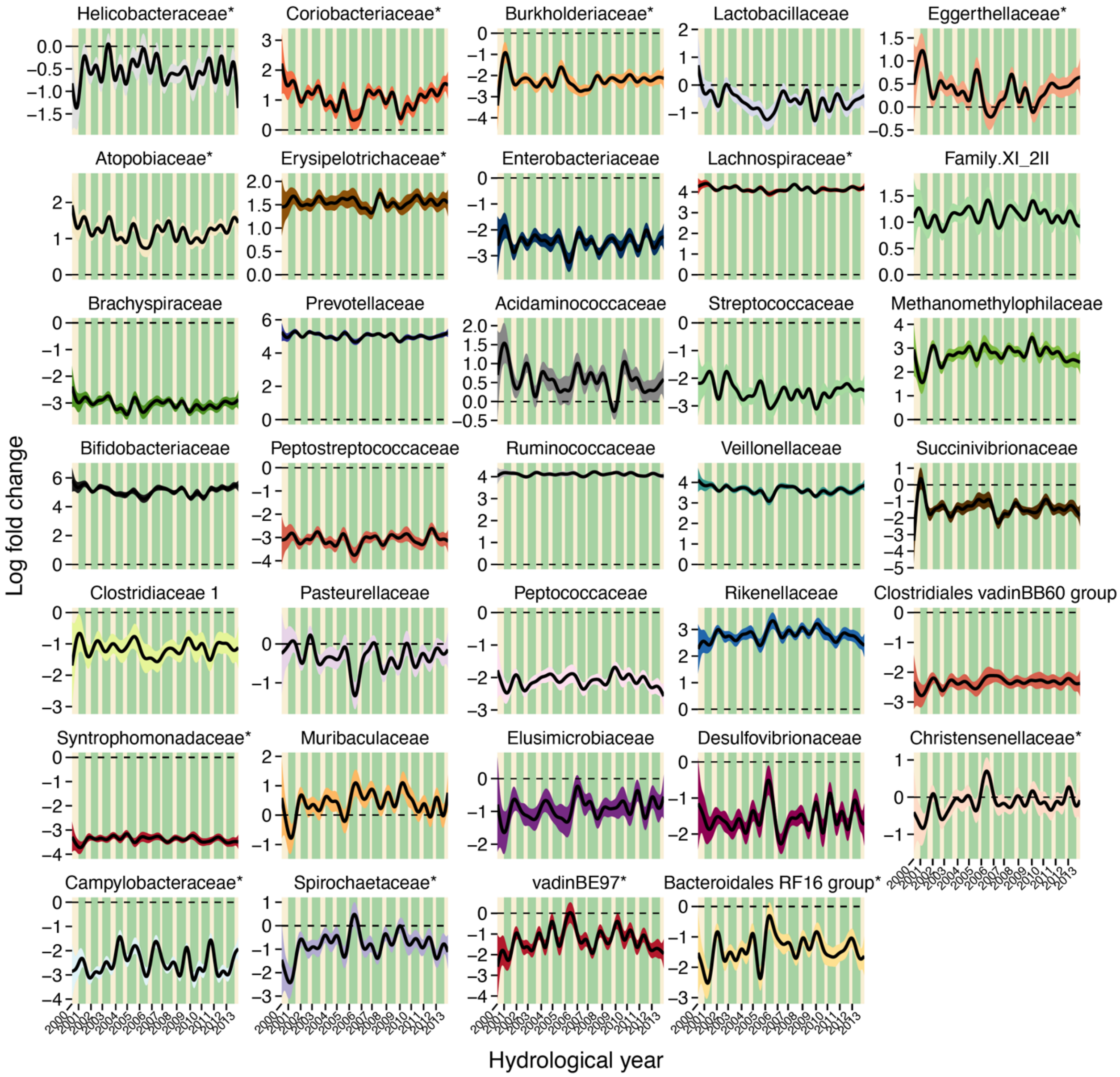


**Fig. S9.** Change in clr-transformed abundances across seasons and years for 34 bacterial families present in >20% of samples. Families are presented in the order they appear in Fig. 2C, starting with those that have the largest log fold increase in the wet season and ending with those that have the largest log fold increase in the dry season. Changes in the clr-transformed relative abundances are expressed as log fold changes, which represent changes in the focal taxon's relative abundance compared to changes in the relative abundance of an average taxon in the same microbial community. A positive/negative log fold change corresponds to taxa whose relative abundance is higher/lower than the average taxon in the same microbial community. Black lines represent fitted smooths; colored bands represent 95% confidence intervals. Each year on the x-axis reflects a hydrological year; green panels correspond to the wet season and yellow panels correspond to the dry season. Asterisks denote taxa that were significant (FDR threshold = 0.05 for n = 393 models) in the linear models Fig. 2C and table S2.

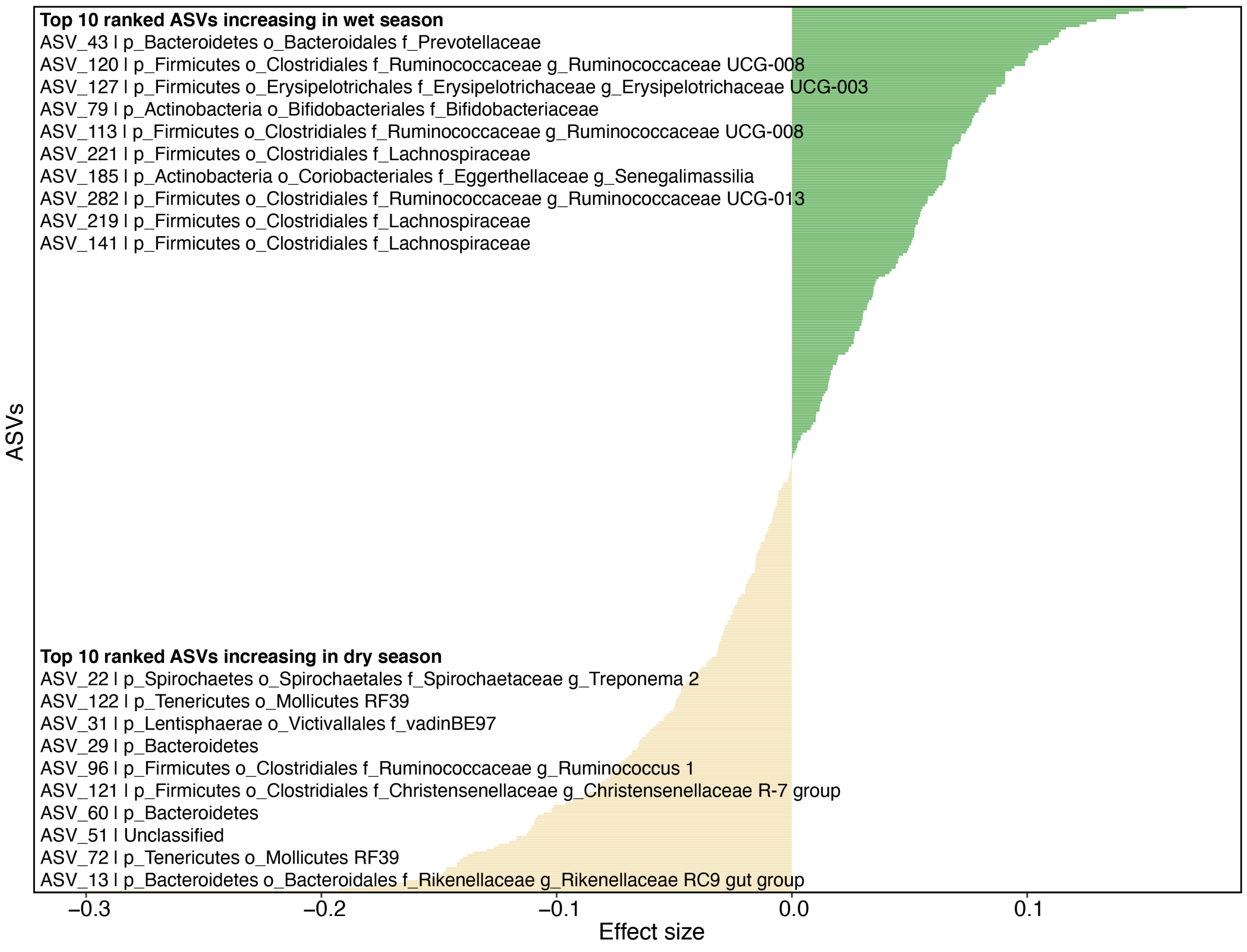


**Fig. S10.** The effect of season varies for the 341 most prevalent ASVs in the data set (i.e., those present in at least 20% of samples). See table S3 for full model results and taxonomic information for each ASV. Horizontal bars show the effect of season from linear models; green bars are ASVs that increase in abundance in the wet season; yellow bars are ASVs that increase in the dry season. The top 10 ASVs (with their taxonomy) with the strongest changes in each season are listed in each corner. Samples from the same host in the same social group collected the same date were averaged prior to running the linear models.

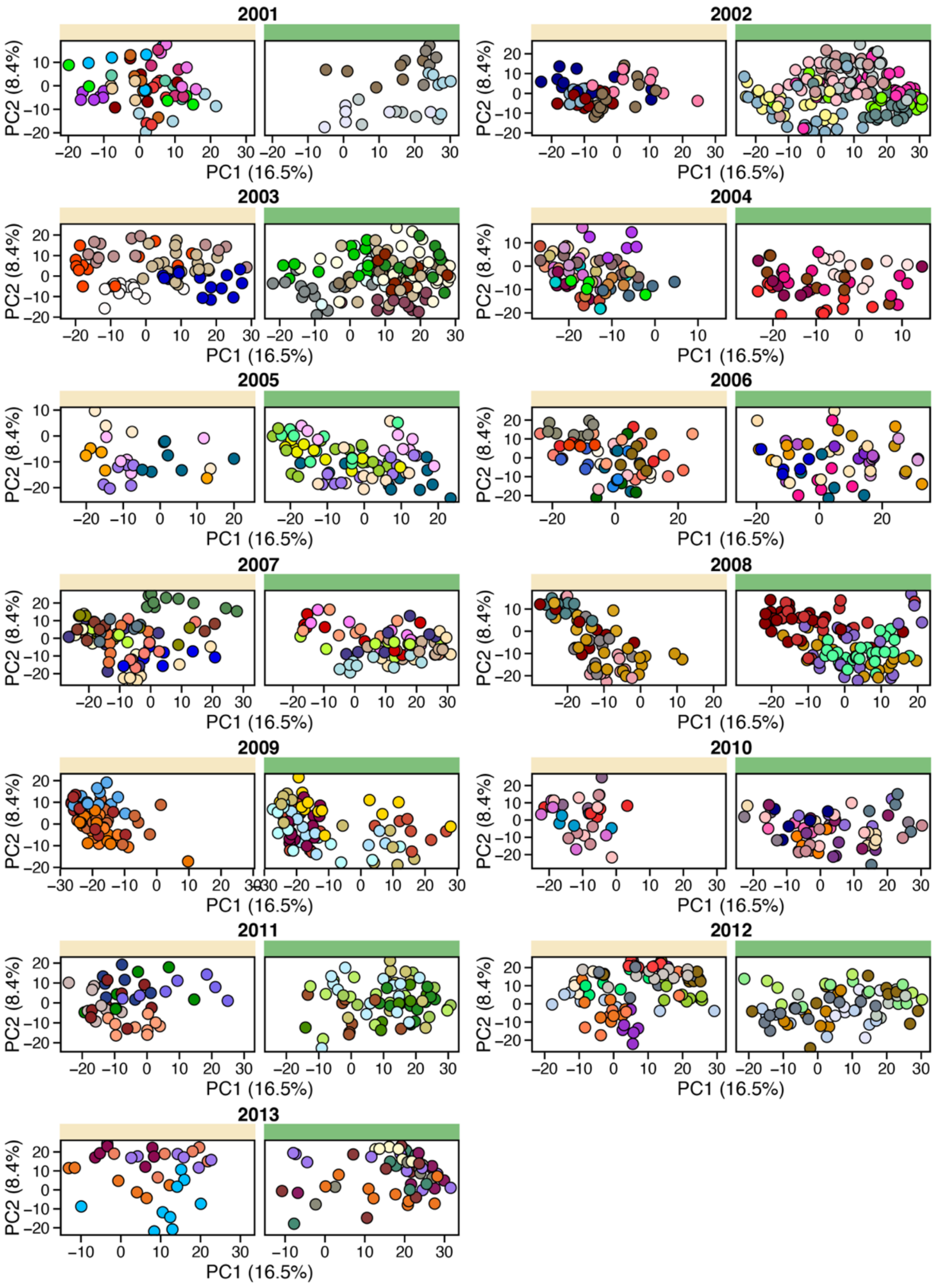


**Fig. S11.** Ordination plots depicting microbiome PC1 and PC2 for the 5 hosts with the most samples in each season and hydrological year (see **fig. S3** for ordinations of all samples; if individuals were tied for the number of samples, more than 5 hosts are depicted). Each point represents an individual sample, colors represent different hosts, and the facet colors correspond to dry (yellow) and wet (green) seasons, respectively. The color of hosts is only preserved within the same year.

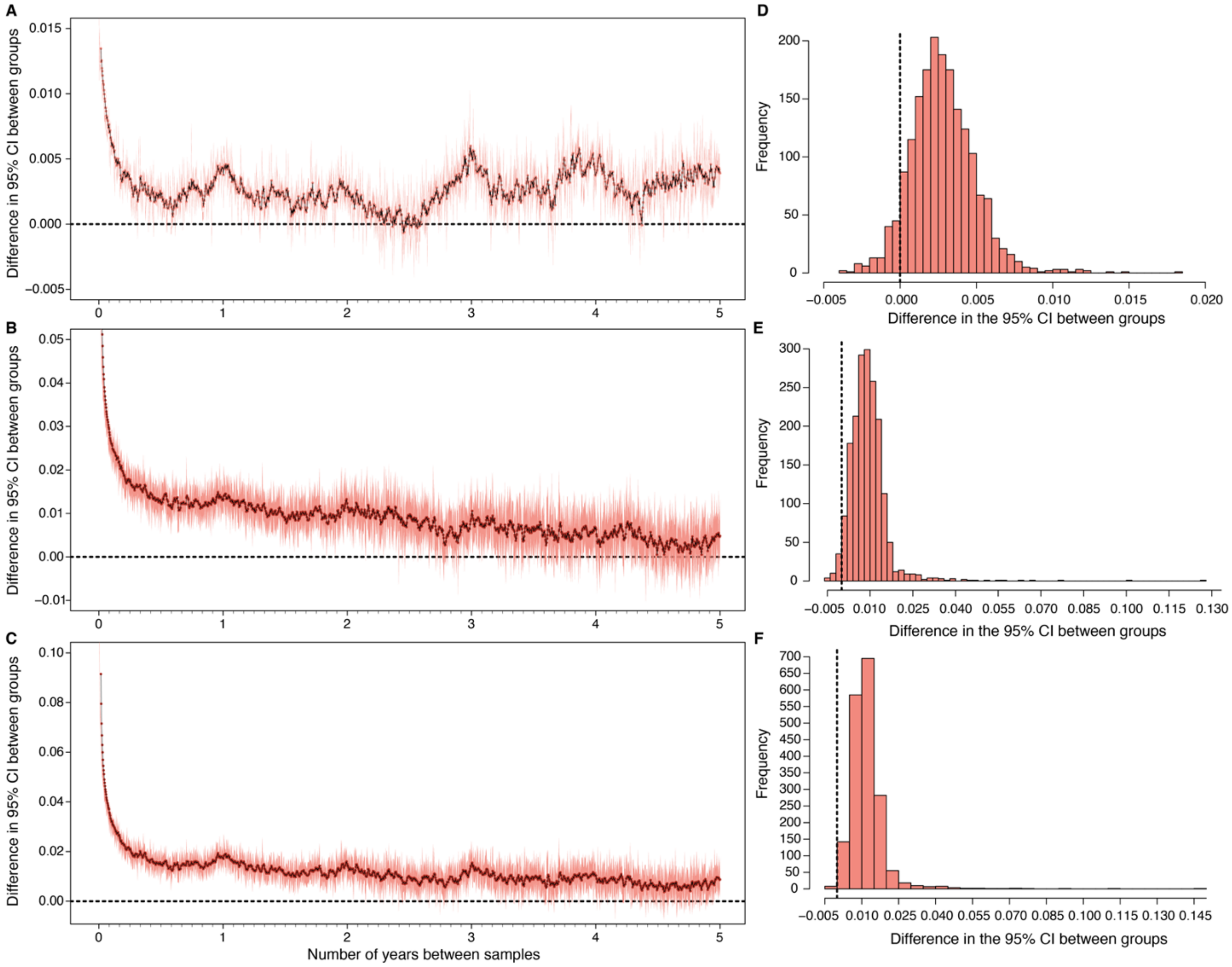


**Fig. S12.** Difference in the 95% confidence intervals (CI) for the three categories of host pairs shown in Fig. 3A in the main text. **(A)** shows the difference in CI between samples collected from different hosts living in the same and different social groups (green vs orange line in **Fig. 3A**) over time; **(B)** shows the difference in CI between samples collected from the same and different hosts in the same social group (brown vs green line in **Fig. 3A**) over time; **(C)** shows the difference in CI for samples from the same host in the same social group and different hosts in different social groups (brown vs orange line in **Fig. 3A**) over time. Dark red lines and lighter red ribbons represent the difference between their moving averages and the 95% CI. Panels **(D-F)** show the corresponding histogram of the differences shown in corresponding panels A-C, with the black vertical dashed line indicating zero difference.

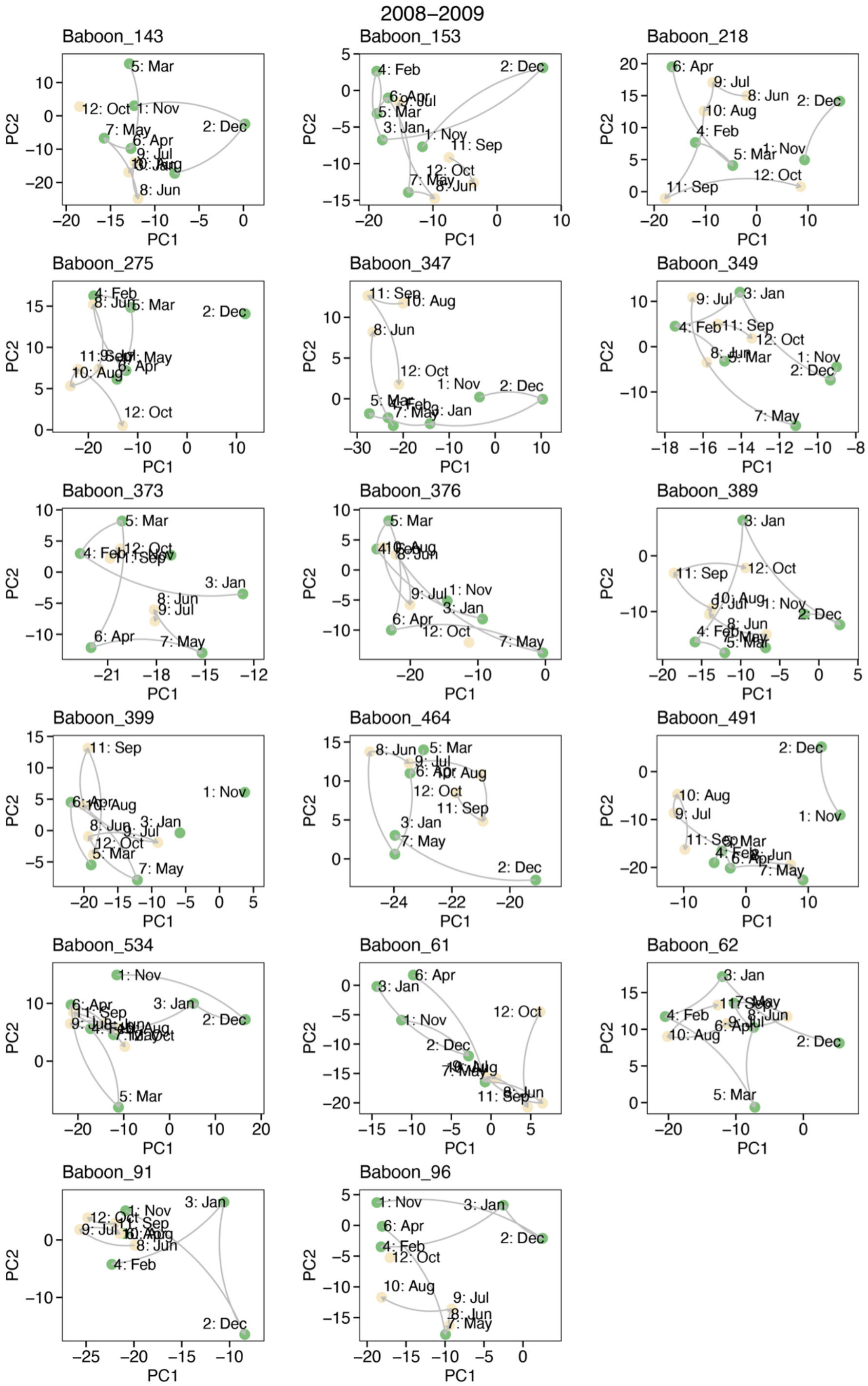


**Fig. S13.** Idiosyncratic microbiome dynamics represented by paths in ordination space. During the 2008-2009 hydrological year, we collected one or more samples for 17 baboons during 10 of the 12 months in that year (N = 174 samples from 17 hosts). Over this time period, these 17 hosts took different paths over the ordination space over this same 1-year span. Each facet represents an individual baboon, each data point represents a sample, labeled with its collection month. Colors show whether the sample was collected in the wet (green) or dry (yellow) season. Grey lines connect samples that are adjacent in time.

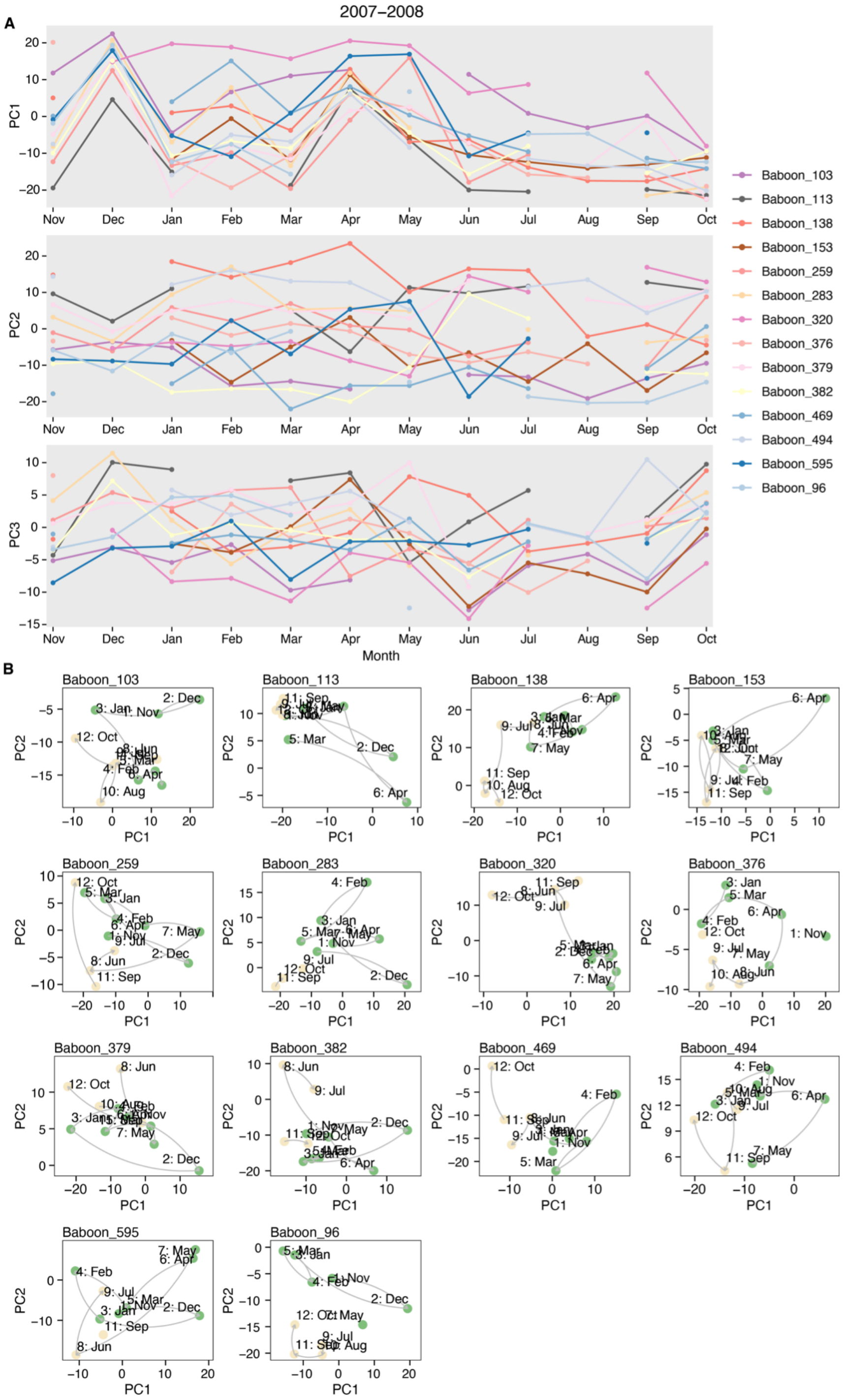


**Fig. S14.** Microbiome dynamics for 14 baboons for which we had at least one sample collected during 10 of the 12 months of the 2007-2008 hydrological year (Nov 2007 to Oct 2008; N = 145 samples from 14 hosts). **(A)** The top three panels show each individual's values for microbiome PC1, PC2, and PC3, and each colored line represents a distinct host. Hosts exhibit widely divergent values of the top 3 principal components of microbiome composition and **(B)** consequently took different paths over the ordination space, shown in the facet plots. See fig. S12 for similar results during the 2008-2009 hydrological year.

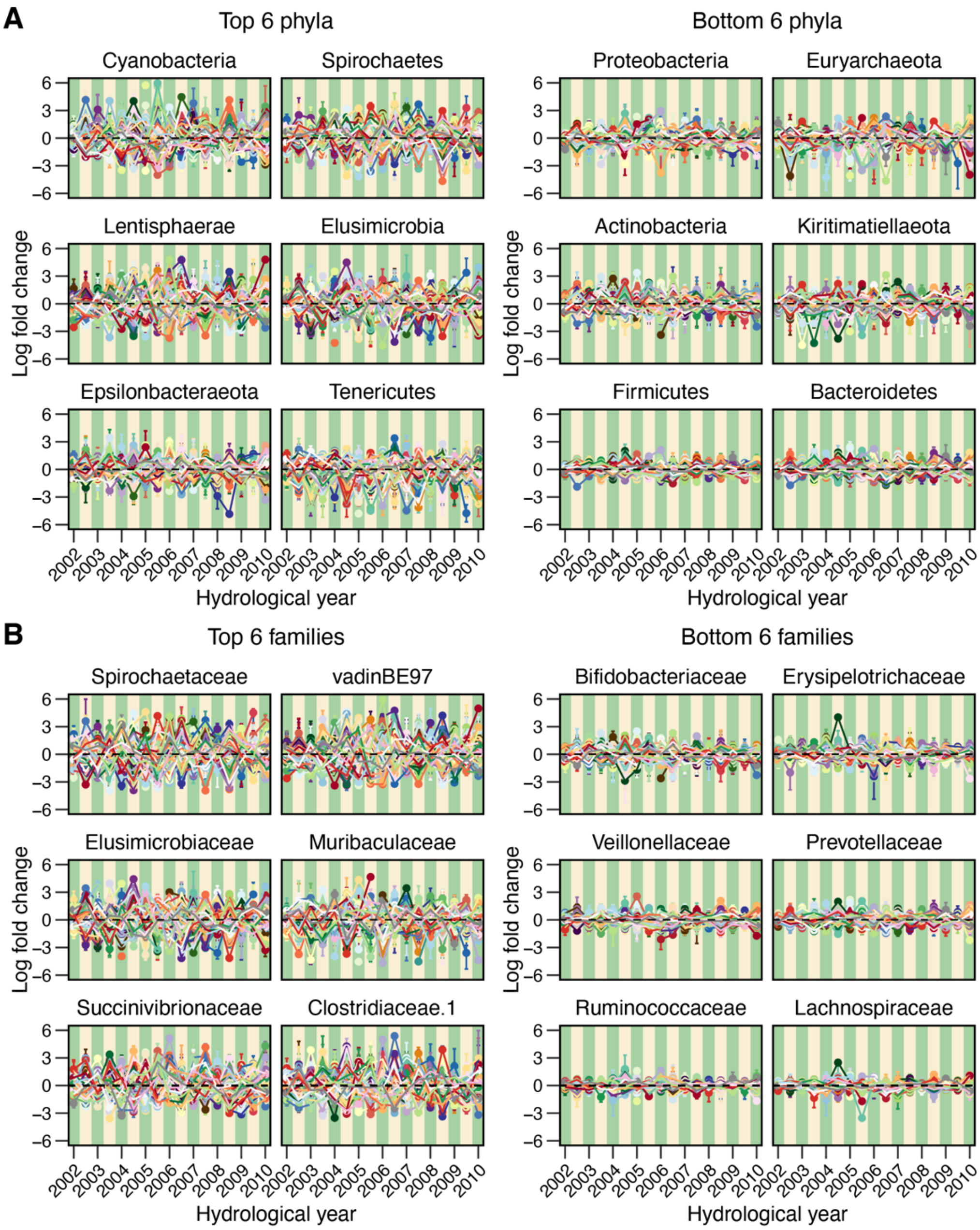


**Fig. S15.** Plots showing variation in log fold changes in microbiome **(A)** phyla and **(B)** families that vary the most (left-hand panels) and least (right-hand panels) between the 56 best-sampled baboon hosts between 2002 and 2010 (see **fig. S2**). See Fig. 3E for visualizations of between-host variation in abundance for all phyla and families. Each point depicts the deviation of a given taxon's average clr-transformed relative abundance in any given host animal, hydrological year and season to its average clr-transformed relative abundance in the host population at large in the same hydrological year and season. A positive/negative log ratio value corresponds to taxa whose relative abundance is, on average, higher/lower in the focal host, hydrological year and season compared to in the host population at large in the same hydrological year and season. Green and yellow background stripes correspond to the wet season and dry season, respectively.

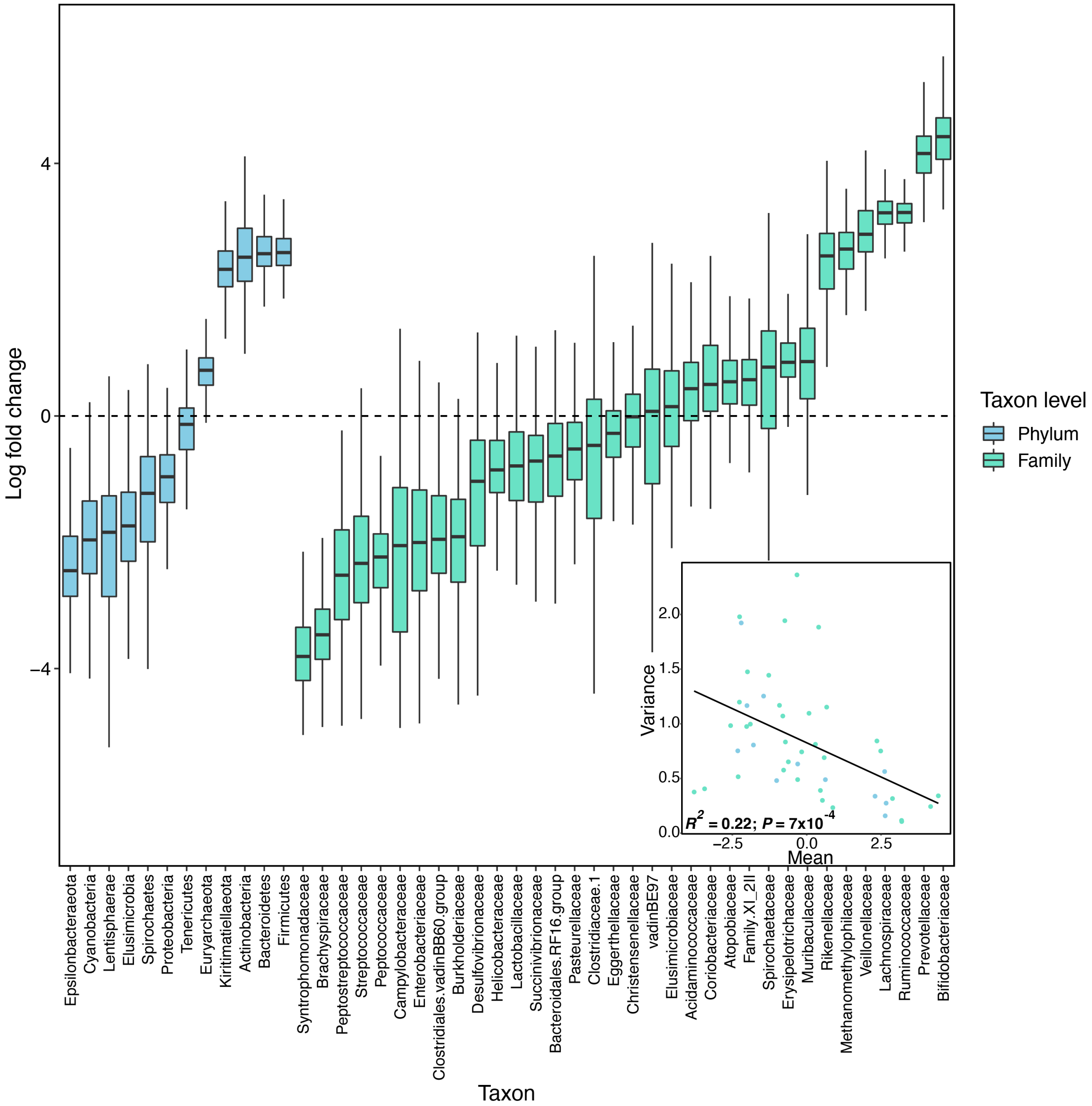


**Fig. S16.** Taxa that are highly variable in abundance between hosts tended to exhibit, on average, below-average abundance compared to less variable taxa. **(A)** box plots showing the distribution of each taxon's clr-transformed relative abundance across the 600 baboon hosts. The y-axis shows the log fold change for each taxon (x-axis). Blue and turquoise box plots represent phyla and families, respectively. Box plots are ordered by their median clr-transformed relative abundance. Taxa with a median above the dashed zero-line have an above-average abundance in more than 50% of the hosts. **(B)** The relationship between the variance in mean clr-transformed relative abundance (y-axis) and the mean clr-transformed relative abundance across hosts (x-axis). Each point represents a taxon, with blue and turquoise dots depicting phyla and families, respectively.

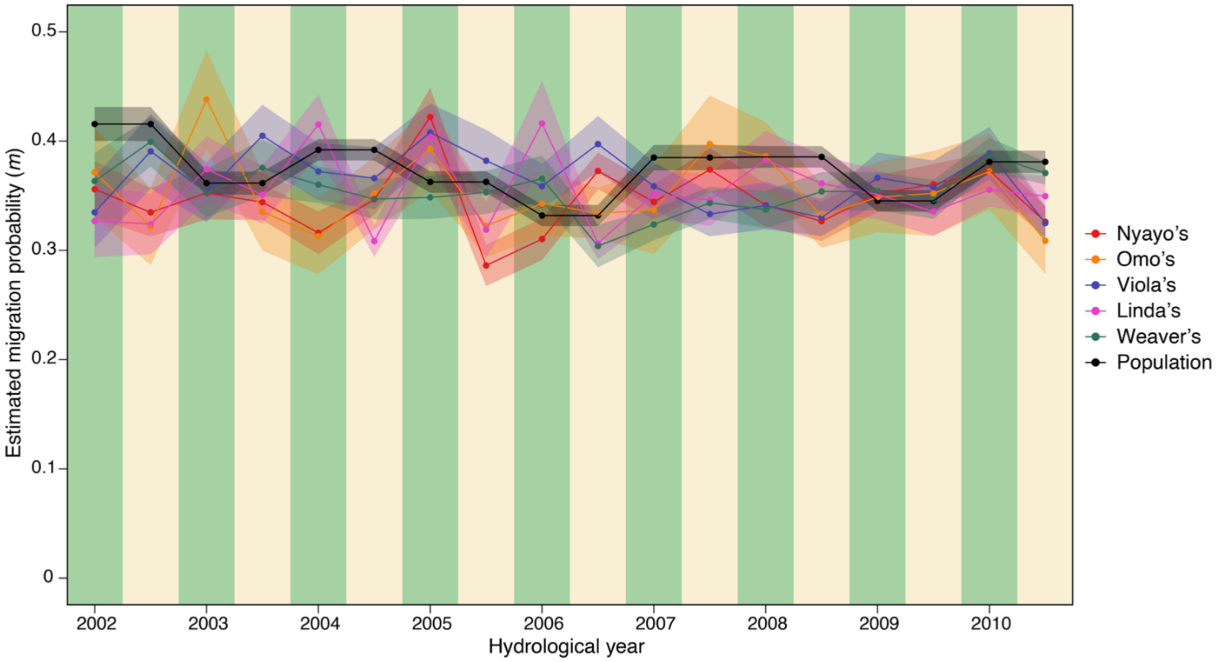


**Fig. S17.** Estimates of metacommunity-wide microbial migration probabilities ($m$) from the Sloan Neutral Community Model for Prokaryotes for ASVs across the whole population (black points and line) and in each social group (colored points and line) in each season and year. The average population-wide estimate of $m$ was 0.373 (range = 0.332 to 0.416). This migration probability was similarly high if we defined the metacommunity to be the host's social group (colored points; average $m$ across groups = 0.355; range = 0.347 to 0.365). Hence, social groups do not seem to represent large barriers to microbial colonization between baboon hosts in different social groups.

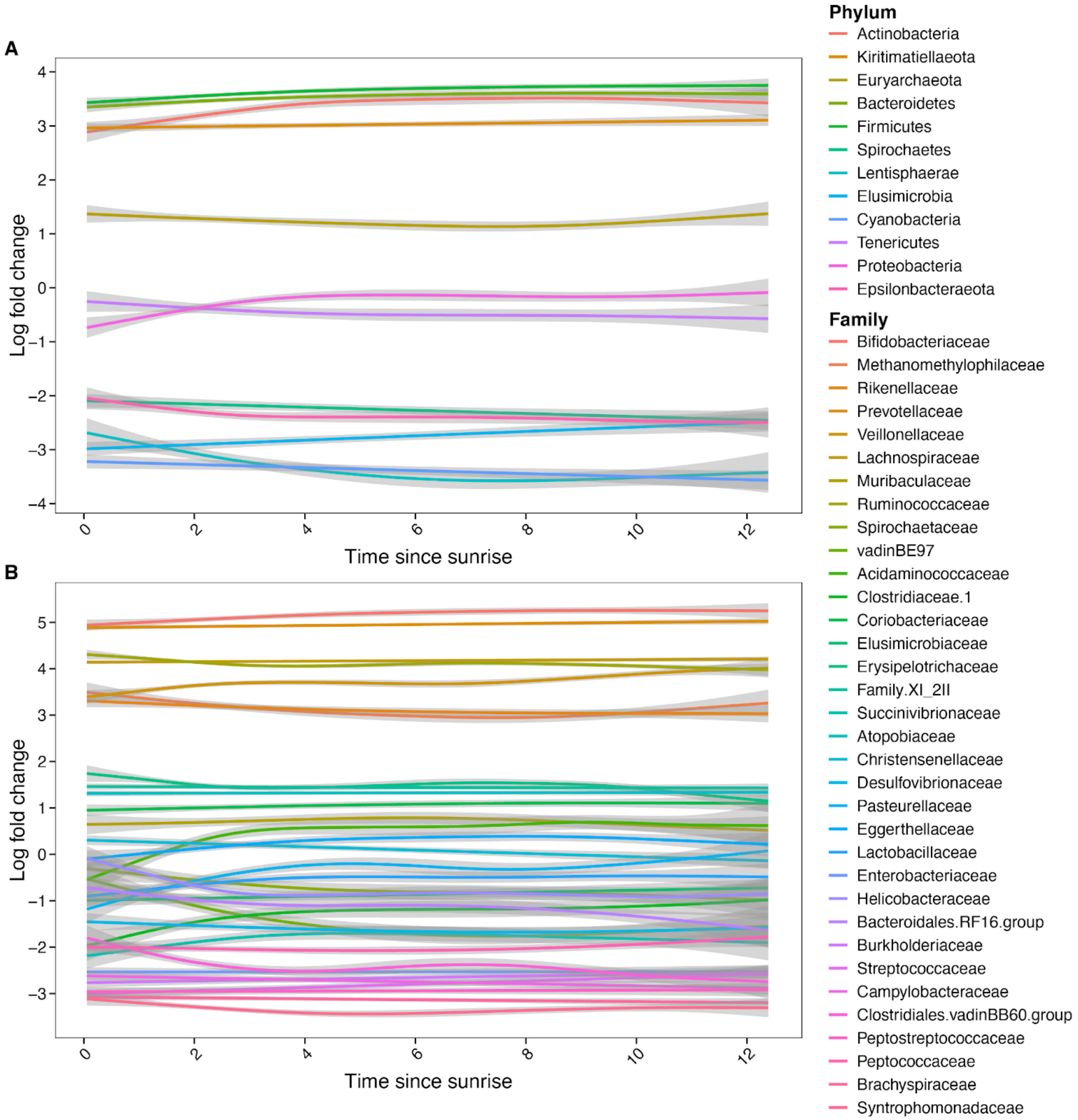


**Fig. S18.** Changes in log fold abundance in **(A)** 12 phyla and **(B)** 34 families as a function of sample collection time since sunrise (in hours). Data on sample collection time were available for 11,517 of the 17,265 samples in our data set. We also repeated the PERMANOVA in **Table S4**, including time since sunrise as the second variable (between day and month of sample collection). Time since sunrise only explained 0.063% of the variation in Aitchison distances and was among the weakest variables in the model.

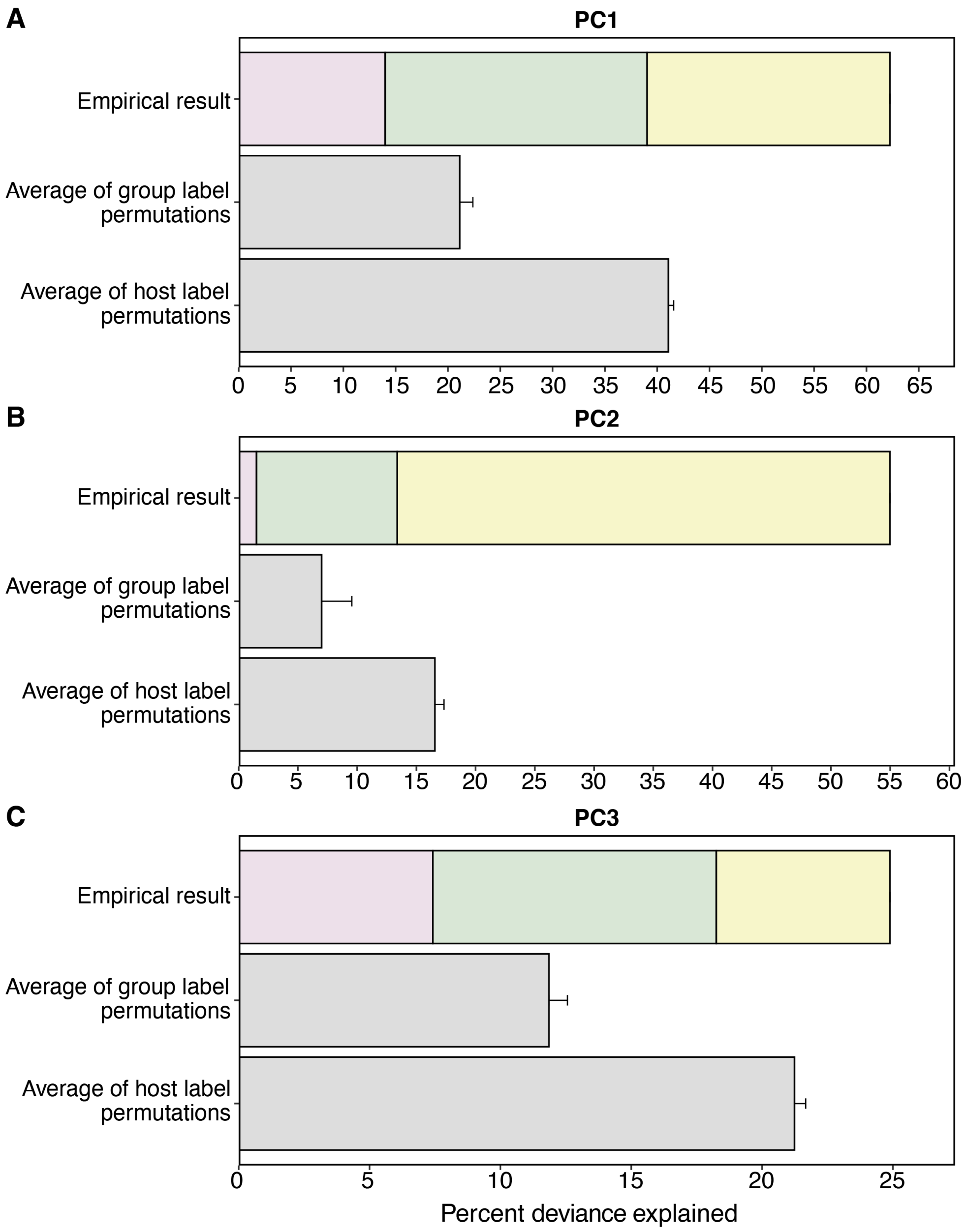


**Fig. S19.** Bar plots showing percent deviance explained by GAMs for **(A)** PC1, **(B)** PC2 and **(C)** PC3. The first bar in each panel ('empirical result') shows the deviance explained by model P, model P+G and model P+G+H as shown in Fig. 4B in the main text. The second bar ('Average of group label permutations') shows the average from 10 permutations of model P+G+H for which we randomized social group identity and group-level covariates across samples while keeping each sample's host identity intact. The third bar ('Average of host label permutations') shows the average from 10 permutations for which we randomized host identity and traits across samples while keeping each sample's annual, seasonal, and group identity intact.

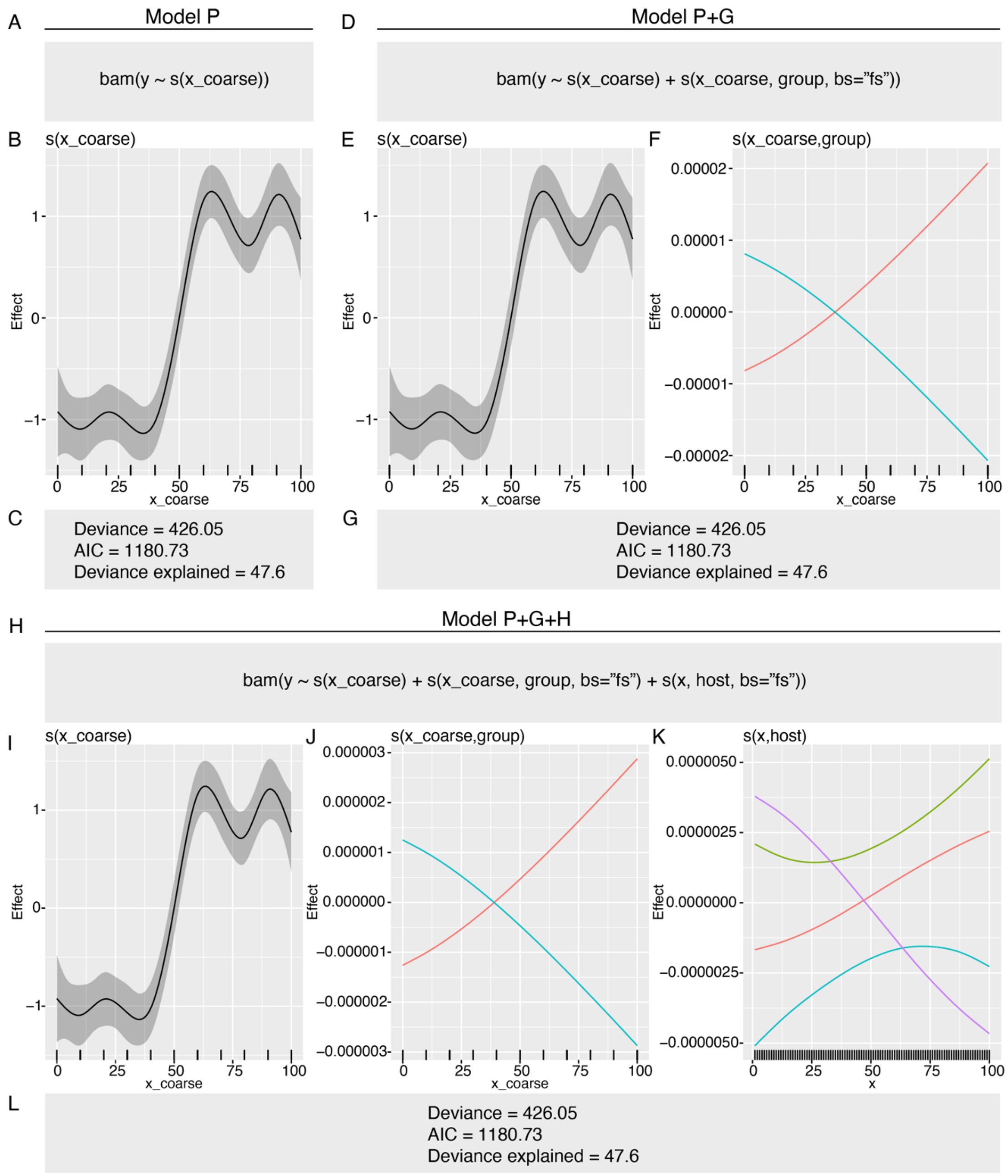


**Fig. S20.** Result from the GAM simulation. Panel **(A-C)** show the results for model P run on the simulated data set. Panel **(D-G)** show the results for model P+G run on the simulated data set. Panel **(H-L)** show the results for model P+G+H run on the simulated data set. All estimated smooths (Panels B, E, F, I, J and K) are drawn by the R package gratia. These plots show the effect of the predictor variable (x) on the response (y) as a function of the predictor variable (x). This simulation experiments show that goodness-of-fit measures (deviance, AIC, deviance explained) are identical for model P+G+H as for model P+G and model P **(C, G and L).** This is because the effect of the host-level and group-level smooth are negligible to model fit.

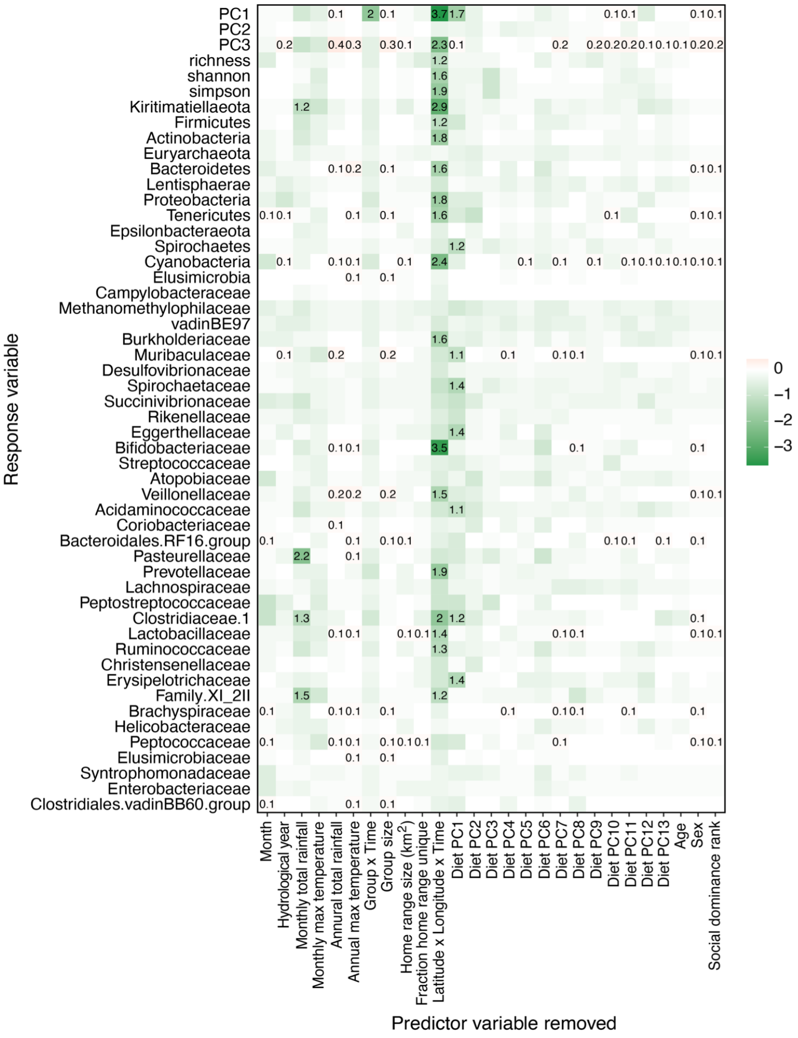


**Fig. S21.** Heat map as in Fig. 4C in the main text showing the loss in percent deviance explained for model P+G+H as we removed each factor in turn from model P+G+H. Data are identical to Fig. 4C, but the variable host x time is not shown to help visualize the effects of other variables in the model. The color in each cell depicts the change in percent deviance explained by removing the variable, with darker shades of green representing greater losses in deviance explained. Numbers in cells indicate the change in deviance from removing a variable from the model. Losses in deviance are shown in green, and we only provide numeric values for losses in deviance > 1%. Gains in deviance are shown in red; we only show numeric values for gains > 0.1%.

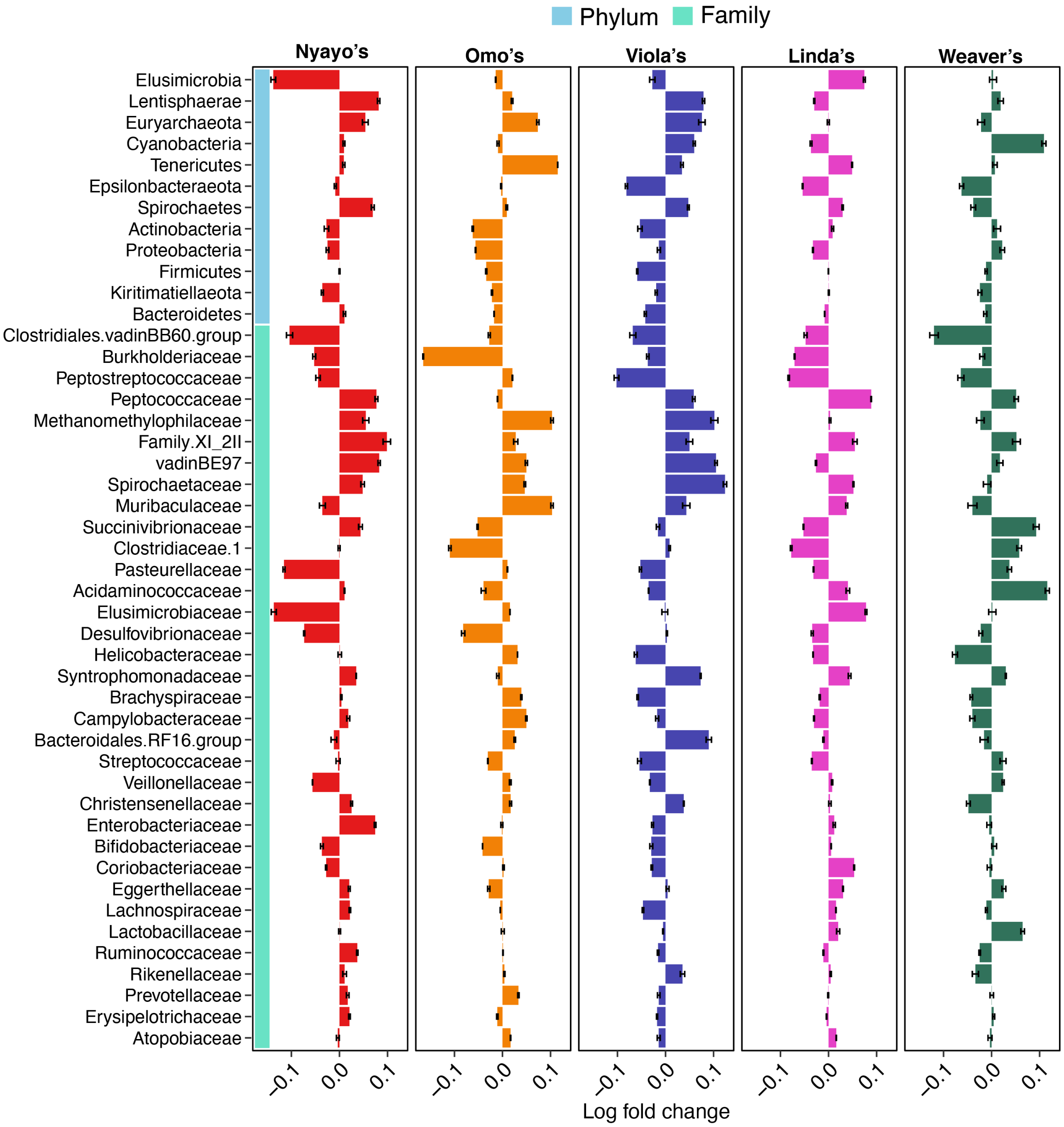


**Fig. S22.** Microbiome phyla and families vary in their contributions to distinctive social group-level compositions. The y-axis shows each of the 12 phyla (light blue vertical bar) and 34 microbiome families (turquoise vertical bar). Each bar depicts the deviation of a given taxon's average clr-transformed relative abundance in each of the 5 original social groups to their average clr-transformed relative abundance in the host population at large. A positive/negative log ratio value corresponds to taxa whose relative abundance is, on average, higher/lower in the focal social group compared to in the host population at large. Taxa are ordered by the sum of their absolute log fold change across social groups.

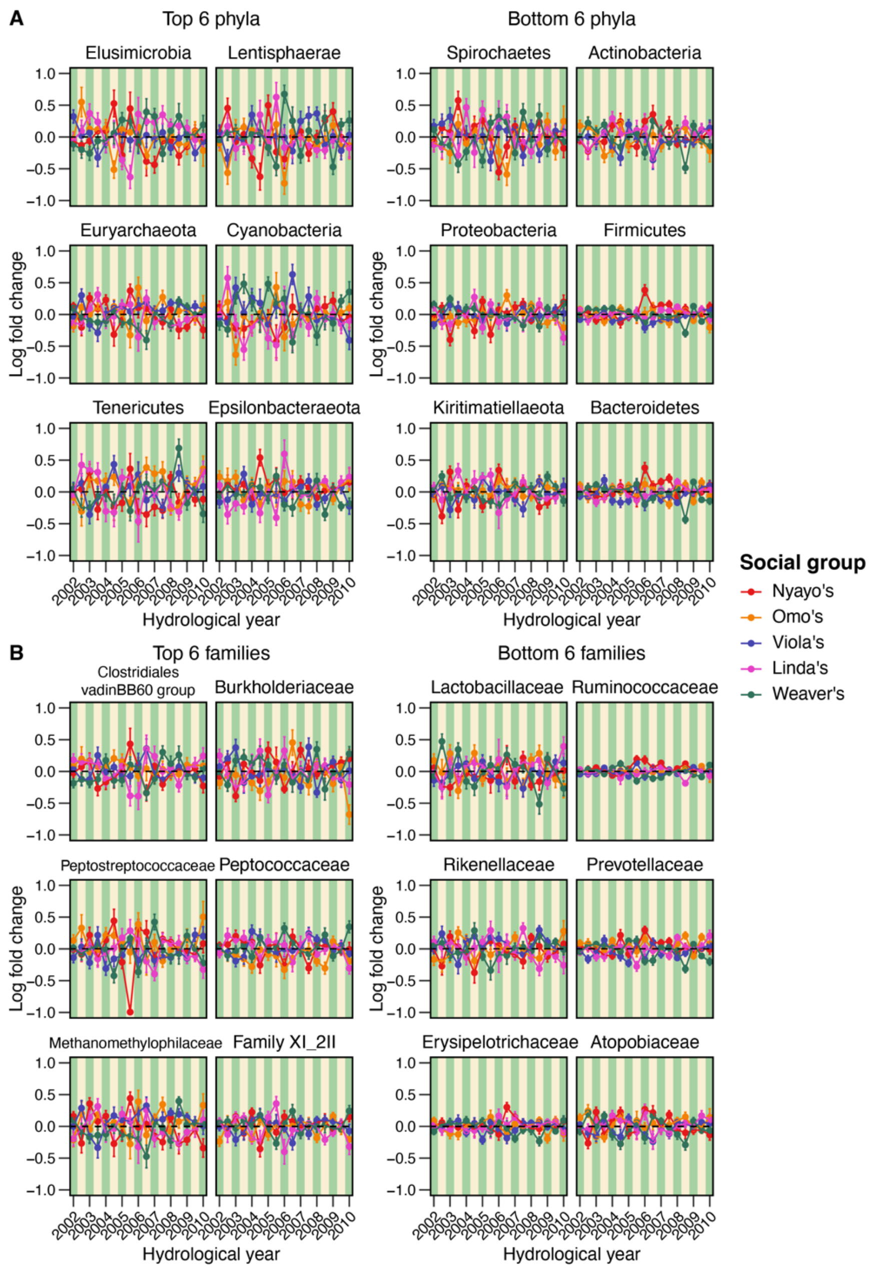


**Fig. S23.** Plots showing variation in log fold changes in microbiome **(A)** phyla and **(B)** families that vary the most (left-hand panels) and least (right-hand panels) between the 5 primary baboon social groups (colors as in Fig. 1). Each point depicts the deviation of a given taxon's average clr-transformed relative abundance in any given social group, hydrological year and season to its average clr-transformed relative abundance in the host population at large in the same hydrological year and season. A positive/negative log ratio value corresponds to taxa whose relative abundance is, on average, higher/lower in the focal social group, hydrological year and season compared to in the host population at large in the same hydrological year and season. Green and yellow background stripes correspond to the wet season and dry season, respectively.

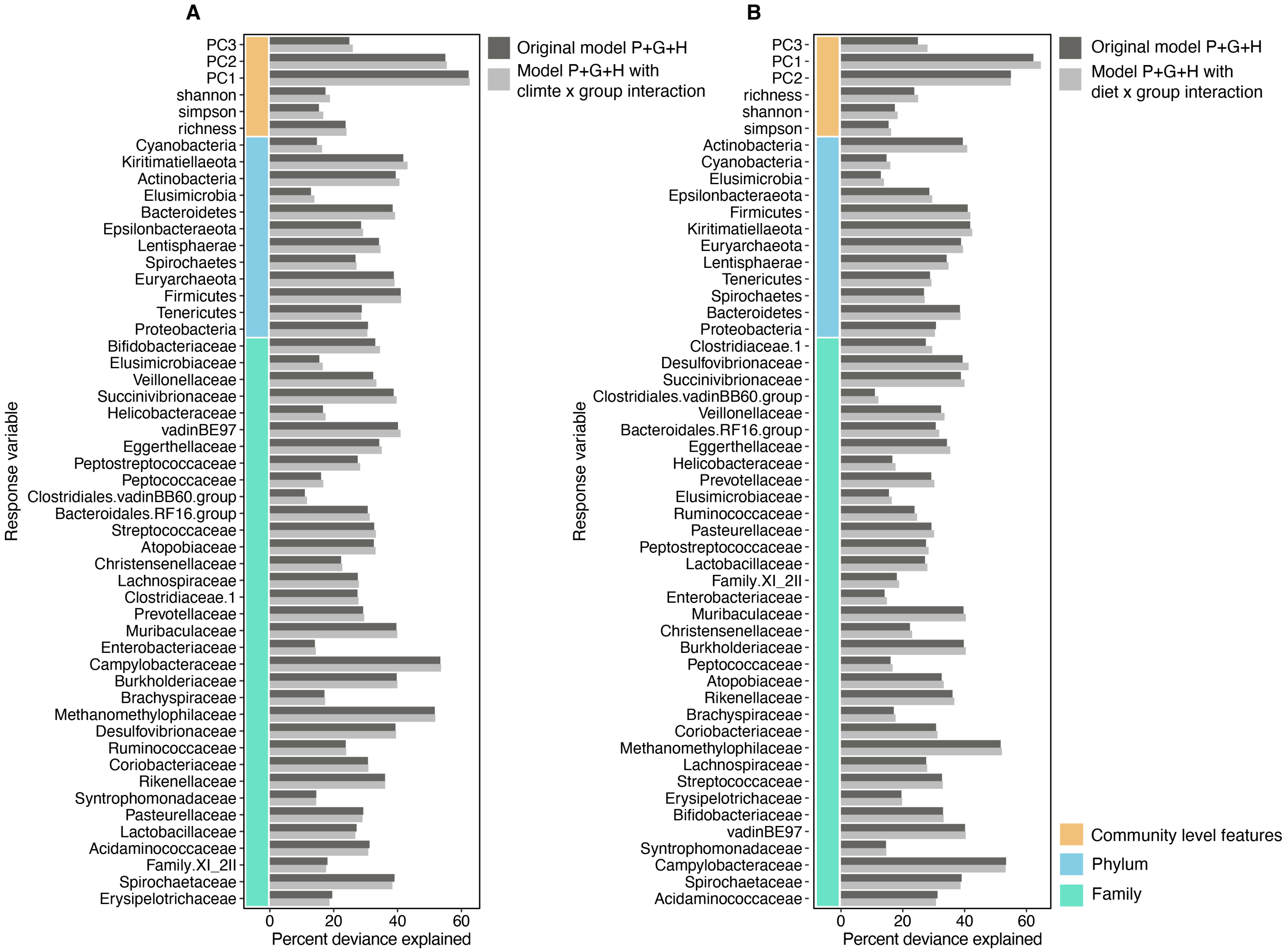


**Fig. S24.** Change in percent deviance explained for versions of model P+G+H that do (dark bars) and do not (light bars) contain interaction effects between **(A)** climate variables and social group identity, and between **(B)** the top three principal components of diet variation and social group identity. The y-axis shows the 52 microbiome features ordered by microbiome feature category (community level features in orange; phyla in light blue; and family in turquoise) and by whether the model without the interactions explains more deviance than the model with the interactions.

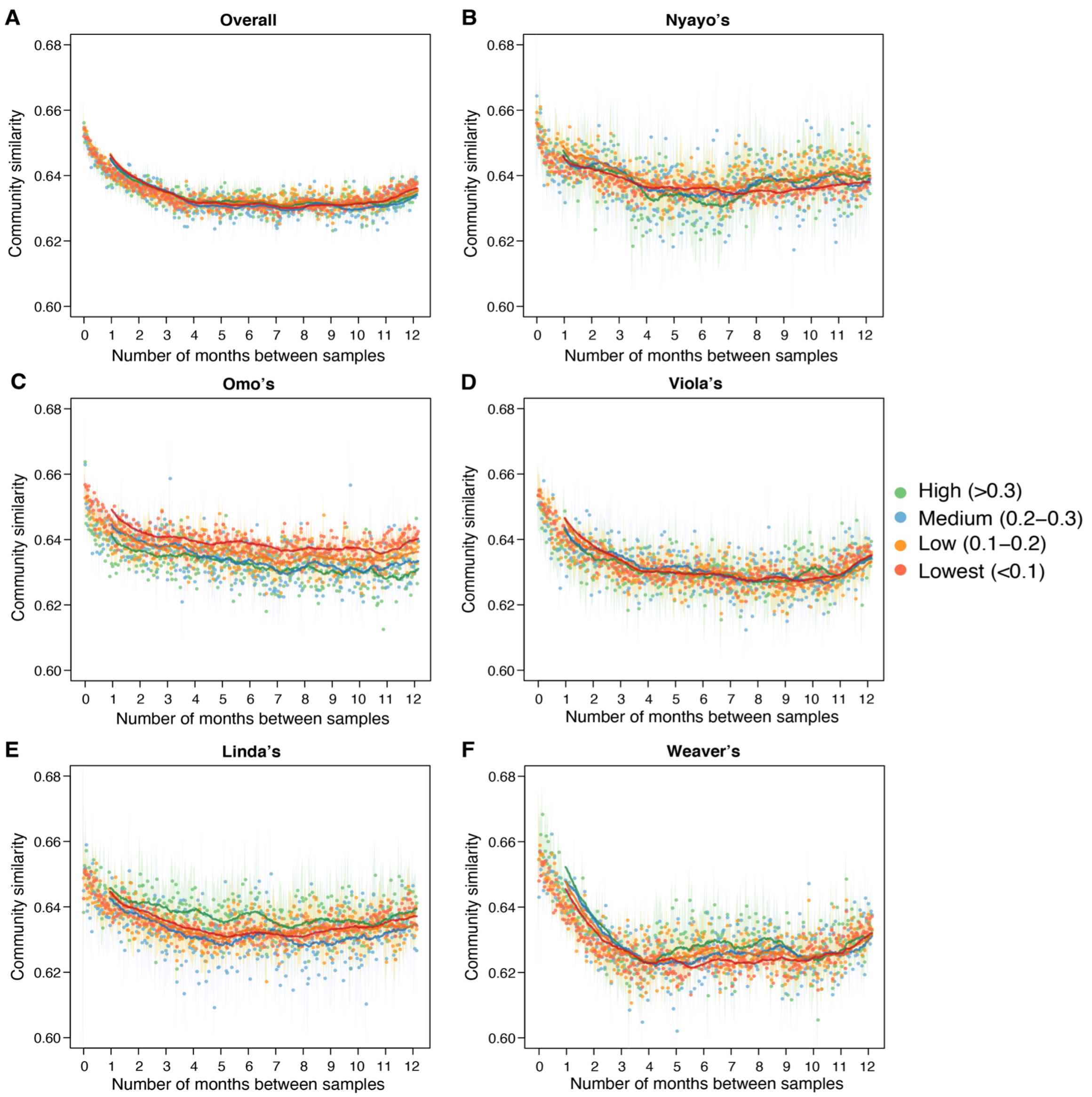


**Fig. S25.** Microbiome composition was not substantially more similar for pairs of baboons who were close grooming partners (green points) versus pairs of baboons who rarely groomed each other (red points). **(A)** Shows results for all social groups combined, while **(B-F)** show results for each of the 5 primary social groups. Grooming relationships reflect all observed instances of grooming between a pair of animals in the past year. Plots show temporal autocorrelation in microbiome Aitchison similarity as a function of the number of days between samples. Green points are pairs of baboons whose grooming relationships were in the top 30% for all grooming pairs in their social group; blue points were in the top 20% to 30% of grooming pairs, orange points were in the top 10% to 20% of grooming pairs, and red points were in the bottom 10% of pairs. Grooming interactions were normalized by the maximum value in each adjacency matrix, and hosts below the age of 5 were removed from the analysis.

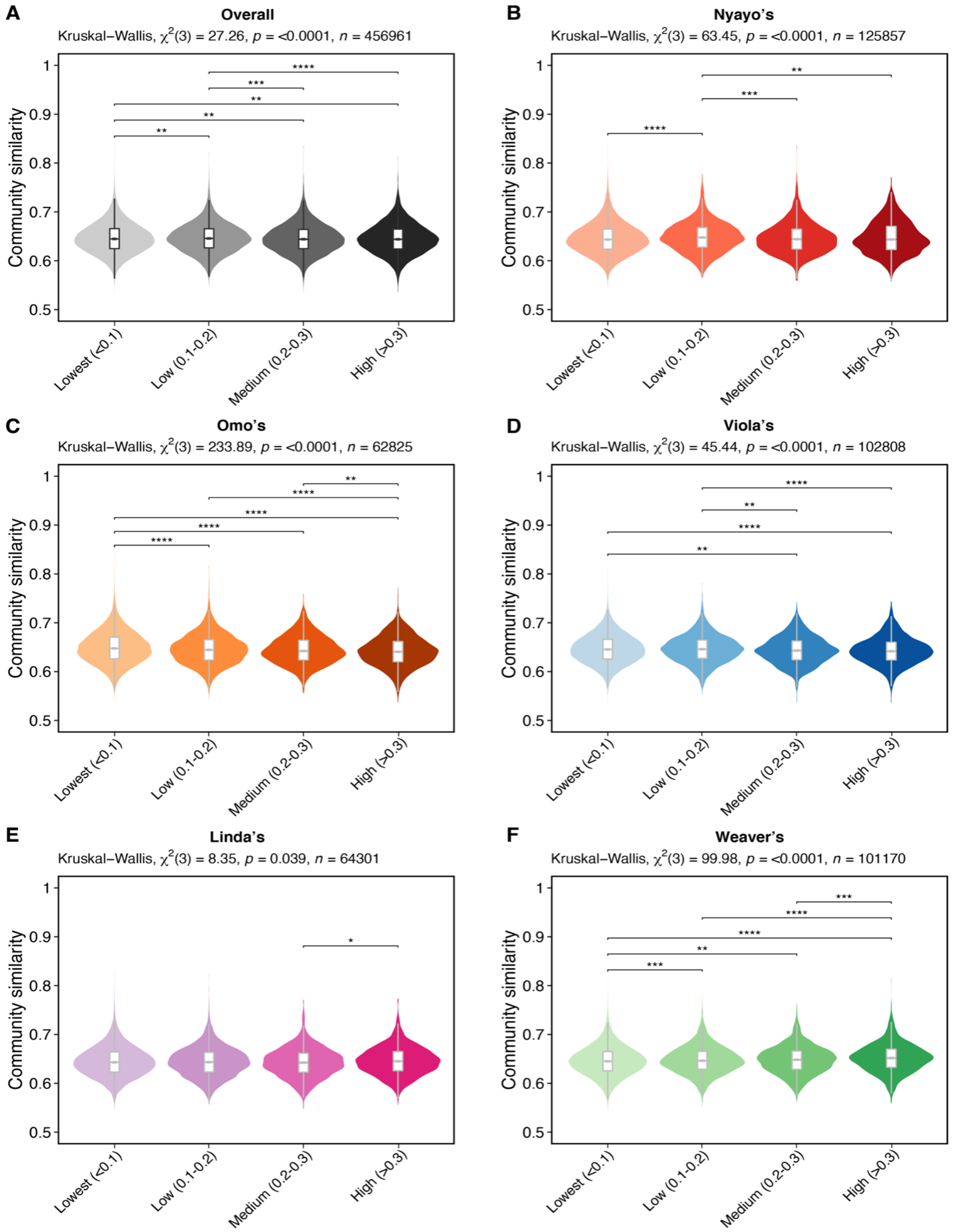


**Fig. S26.** Average Aitchison similarity among samples collected within 30 days of each other across social groups **(A)** and within each of the 5 main social groups **(B-F)**. Within each plot, the gradient (from left to right) corresponds to hosts with lowest (<0.1); low (0.1-0.2); medium (0.2-0.3); and high (>0.3) grooming interaction with other hosts in the same social group. Grooming interactions were normalized by the maximum value in each adjacency matrix, and hosts below the age of 5 were removed from the analysis. The Kruskal-Wallis test was used to test for overall significance, and Dunn's Multiple Comparison Test was used for pairwise significance tests with Benjamini-Hochberg FDR adjusted p-values. *: p <= 0.05; **: p <= 0.01; ***: p <= 0.001; ****: p <= 0.0001.

#### B. Supplementary video legends

**Video S1.** Animation of home range use by the 5 original social groups, as defined in Fig. 1. This animation shows the geographical location of each social group over time, with the x-axis showing longitude and the y-axis latitude. Each dot represents the average monthly longitude and latitude per social group and month. While solid dots represent the groups' current position in the focal month and hydrological year, the hollow circles show previous locations, thus outlining each social group's total home range area over time.

**Video S2.** Animation of the microbiome PC1 and PC2 for baboons living in the 5 original social groups, as defined in Fig. 1. To make this animation, we first averaged the PC1 and PC2 sample scores by host and collection date, such that each host only has one value per collection date. We then 'filled in' missing collection dates, and performed a 30-day sliding window analysis (step size=1). Each frame (i.e., image) in the animation corresponds to one sliding window. The date in the top left corner corresponds to the first date of each window, and each dot represents one individual host colored by its social group.
